# Comparative Multi-omic Mapping of Human Pancreatic Islet Endoplasmic Reticulum and Cytokine Stress Responses Provides Insights into Type 2 Diabetes Genetics

**DOI:** 10.1101/2024.02.17.580646

**Authors:** Eishani Kumar Sokolowski, Romy Kursawe, Vijay Selvam, Redwan M. Bhuiyan, Asa Thibodeau, Chi Zhao, Cassandra N. Spracklen, Duygu Ucar, Michael L. Stitzel

## Abstract

Endoplasmic reticulum (ER) and inflammatory stress responses are two pathophysiologic factors contributing to islet dysfunction and failure in Type 2 Diabetes (T2D). However, how human islet cells respond to these stressors and whether T2D-associated genetic variants modulate these responses is unknown. To fill this knowledge gap, we profiled transcriptional (RNA-seq) and epigenetic (ATAC-seq) remodeling in human islets exposed to *ex vivo* ER (thapsigargin) or inflammatory (IL-1β+IFN-γ) stress. 5,427 genes (∼32%) were associated with stress responses; most were stressor-specific, including upregulation of genes mediating unfolded protein response (e.g. *DDIT3, ATF4*) and NFKB signaling (e.g. *NFKB1, NFKBIA*) in ER stress and cytokine-induced inflammation respectively. Islet single-cell RNA-seq profiling revealed strong but heterogeneous beta cell ER stress responses, including a distinct beta cell subset that highly expressed apoptotic genes. Epigenetic profiling uncovered 14,968 stress-responsive *cis-*regulatory elements (CREs; ∼14%), the majority of which were stressor-specific, and revealed increased accessibility at binding sites of transcription factors that were induced upon stress (e.g. ATF4 for ER stress, IRF8 for cytokine-induced inflammation). Eighty-six stress-responsive CREs overlapped known T2D-associated variants, including 20 residing within CREs that were more accessible upon ER stress. Among these, we linked the rs6917676 T2D risk allele (T) to increased *in vivo* accessibility of an islet ER stress-responsive CRE and allele-specific beta cell nuclear factor binding *in vitro*. We showed that *MAP3K5,* the only ER stress-responsive gene in this locus, promotes beta cell apoptosis. Consistent with its pro-apoptotic and putative diabetogenic roles, *MAP3K5* expression inversely correlated with beta cell abundance in human islets and was induced in beta cells from T2D donors. Together, this study provides new genome-wide insights into human islet stress responses and putative mechanisms of T2D genetic variants.

## INTRODUCTION

Type 2 diabetes (T2D) is a complex metabolic disorder, characterized by an interplay between genetics and environment that leads to pancreatic islet beta cell dysfunction and/or death, and inadequate insulin secretion in response to insulin resistance^1–5^. Genome-wide association studies (GWAS) have linked DNA sequence variants in >600 loci in the human genome with increased T2D risk or progression^6^. The abundance of non-coding locations of these variants, combined with previous studies demonstrating significant enrichment of variants in islet *cis*-regulatory elements (CREs), suggests that these variants contribute to islet dysfunction and failure by altering CRE use or function and effector gene expression^2,4,7–10^. We and others have discovered that a subset of T2D-associated variants alter *in vivo* CRE chromatin accessibility and/or effector gene expression in human islets under steady-state conditions^4,7–9,11–13^. However, as T2D pathogenesis is heavily influenced by the dynamic interaction between genetic variants and environmental stressors^1,2,4,5^, the functional effects of these variants, particularly in the context of islet stress responses such as endoplasmic reticulum (ER) stress and pro-inflammatory cytokine responses, are largely unknown.

ER stress is crucial in the context of T2D as it is integral to protein quality control and insulin synthesis in beta cells^14,15^. Under chronic hyperglycemia, a sustained demand for insulin production can overwhelm the beta cell ER, leading to heightened stress and activation of the unfolded protein response (UPR) machinery^14^. Prolonged or excessive ER stress can contribute significantly to beta cell dysfunction and death^11,14,15^. Beta cell dysfunction has been further linked to high levels of pro-inflammatory cytokines in the blood^15–17^, which have been shown to trigger the NFKB pathway, resulting in impaired insulin secretion^17–19^. Although ER and inflammatory stressors have been associated with T2D^20,21^, it is unclear how pancreatic islet cells respond to each specific stressor and whether any T2D-associated variants are linked to response-associated genomic regions.

To fill these knowledge gaps, we defined transcriptional regulatory programs controlling human islet responses to ER stress and pro-inflammatory cytokines by mapping genome-wide CRE accessibility (via ATAC-seq) and gene expression (via RNA-seq) in islets exposed to the ER stress-inducing agent thapsigargin or inflammation-inducing cytokines (IL-1β and IFN-γ). Comparison of the stress response genes and CREs revealed complementary, stress- and cell type-specific changes in transcriptional regulatory programs and expression of the factors mediating these stress responses. We identify T2D- associated variants in 38 signals overlapping ER stress- or cytokine-induced CREs as candidate causal variants and link them to stress-responsive target (and putative T2D effector) genes. Targeted variant-to-function analyses in the *SLC35D3* locus link the rs6917676 T2D risk allele to increased ER stress-responsive CRE accessibility and demonstrate that the putative T2D effector *MAP3K5*, the only ER stress-responsive gene in the locus, promotes stress-responsive beta cell apoptosis.

## RESULTS

### Comprehensive comparative mapping of ER stress- and cytokine-responsive genes in human islets

To define the characteristic responses of human pancreatic islets to ER stress and pro-inflammatory cytokines, we procured primary human islets from 30 non-diabetic donors **(Supplementary Table 1)** and exposed them to a 24-hour treatment with either thapsigargin (vs. DMSO solvent control)^11,22,23^ or an IL- 1β+IFN-γ cocktail (vs. untreated control)^24^, respectively. We determined and compared the genome-wide gene expression changes elicited by these two T2D-relevant stressors using whole islet RNA sequencing (RNA-seq)^25^.

In total, ∼32% (5,427/17,096) of autosomal protein-coding genes **(Methods)** expressed in human pancreatic islets responded significantly (FDR<5%; |FC|≥1.5) to at least one of the stressors compared to control conditions **(Supplementary Table 2)**. 2,967 genes were differentially expressed (DE) upon ER stress (1,517 induced; 1,450 reduced), whereas 3,443 genes were DE upon cytokine-induced inflammation (1,893 induced; 1,550 reduced) **(Supplementary Table 2)**. Transcriptional responses to ER stress and cytokines were largely distinct. For example, ∼85% of induced genes were stressor-specific, including 1,064 ER stress-specific genes and 1,440 cytokine-specific genes **(Figure 1A; Supplementary Table 2)**. As anticipated, ER stress treatment induced genes facilitating both the homeostatic (e.g., *ATF4, ERN1, EIF2AK3, HERPUD1, HSPA5*) and terminal (e.g., *DDIT3, MAP3K5*) arms of UPR and ER protein processing related pathways **(Figure 1B; Supplementary Table 2)**, which are centrally linked to regulating insulin synthesis, managing ER stress responses, and controlling apoptosis - critical processes for beta cell function and survival^26–32^ **(Figures 1B-C)**.

**Figure 1:**
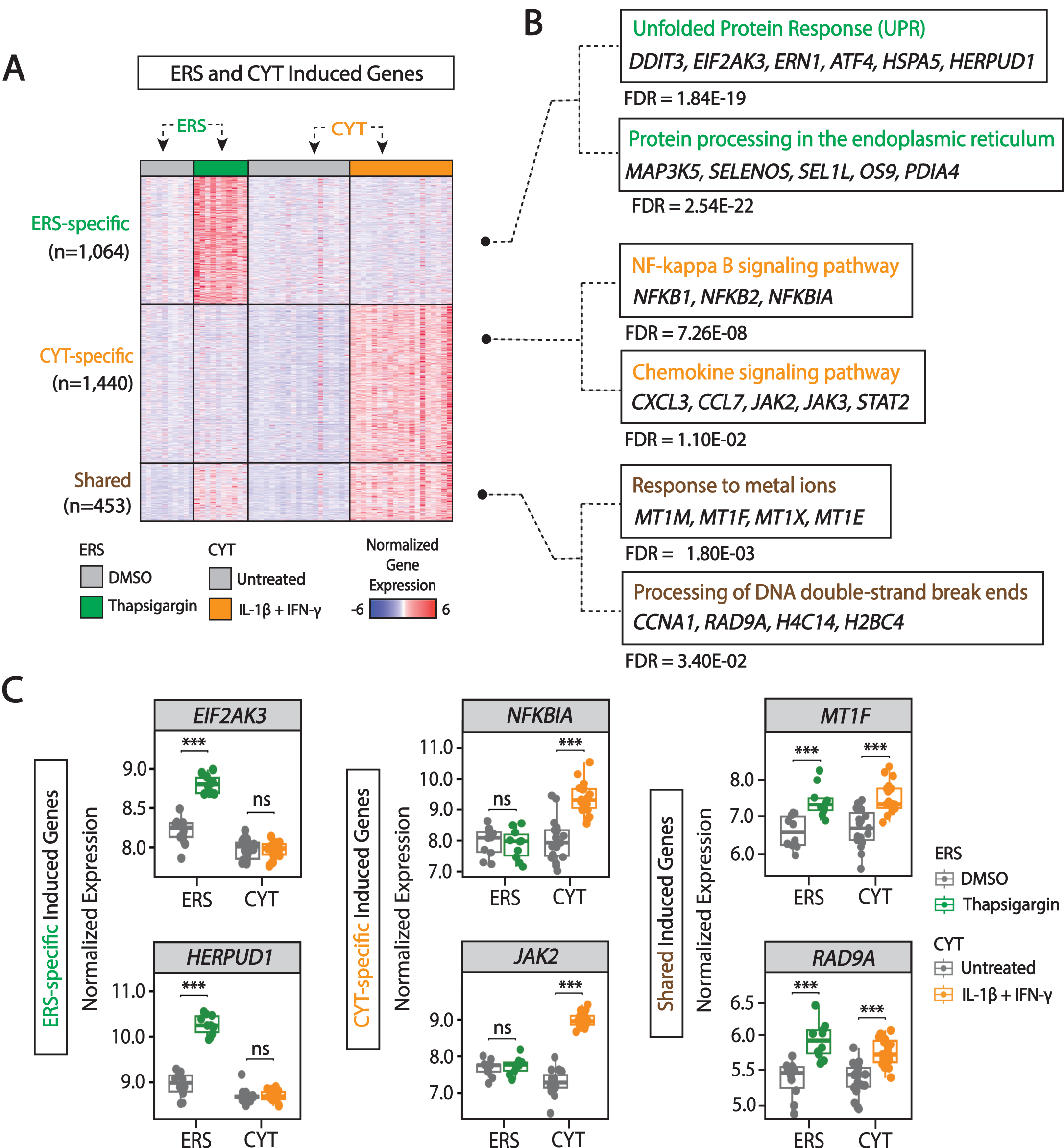
Induced transcriptional responses of human pancreatic islets to ER stress (ERS) and pro-inflammatory cytokines (CYT). (A) Heatmap of genes induced by ERS and/or CYT treatment (FDR<5%; FC≥1.5). Induced genes are categorized as ERS-specific, CYT-specific, or shared between both conditions; the number of genes in each category is denoted in parentheses on the left. Note that the majority of genes exhibit stress-specific induction. Expression values are scaled using z-scores. **(B)** Enriched pathways for induced genes; FDR values and example genes for each pathway are as indicated. **(C)** Examples of enriched pathway genes induced by ERS, CYT, or both. Dot-and-box plots show gene expression levels (CPM) per islet donor in ERS (green), CYT (orange), or control samples (grey). ***=FDR<5% and FC≥1.5; ns=not significant. FDRs were calculated using Benjamini-Hochberg p-value adjustment. FDR, False Discovery Rate; FC, fold change; CPM, counts per million.

Cytokine-induced genes were enriched in NFKB and chemokine signaling related pathways **(Figure 1A; Supplementary Table 2)**, including NFKB complex members (e.g., *NFKB1*, *NFKBIA*) and important signaling molecules (e.g., *JAK2, STAT2*) **(Figures 1B-C; Supplementary Table 2)**, consistent with previous reports^33,34^. These genes have been linked to modulating inflammatory responses, promoting immune cell infiltration, and contributing to islet beta cell dysfunction^35–41^. 453 genes were consistently induced by both ER and pro-inflammatory cytokine stressors **(Figure 1A; Supplementary Table 2)**, which were enriched in pathways related to: 1) metal ion response, including metallothioneins (*MT1* genes), which scavenge free radicals and heavy metals in stressed cells and are associated with reduced insulin secretion upon stress; and 2) processing of DNA double-strand breaks (DSBs), including *RAD9A,* which is involved in repairing DNA damage and double-strand breaks associated with T2D^42–44^ 12/22/23 12:17:00 PM**(Figures 1B-C; Supplementary Table 2)**.

Similarly, ∼79% of reduced genes were stressor-specific, including 920 ER stress-specific genes, and 1,020 cytokine-specific genes **(Supplementary** Figure 1A**; Supplementary Table 2)**. *PDX1, ADCY5, GLP1R* and *IGFBP5*, which encode factors integral to islet identity and function^45–48^ were reduced upon ER stress **(Supplementary** Figures 1B-C**; Supplementary Table 2)**. In contrast, *SLC1A1, COL2A1, NPNT* and *ITGA10*, which participate in protein digestion/absorption and extracellular matrix (ECM) receptor signaling related pathways and are important for beta cell function^49–51^, were reduced upon cytokine-induced inflammation **(Supplementary** Figures 1B-C**; Supplementary Table 2).** 530 genes were reduced by both stressors **(Supplementary** Figure 1A**; Supplementary Table 2)**, including *CDC20, CDC45, UGT2B11* and *UGT2B15*, which are involved in cell cycle and retinol metabolism and are crucial for islet function^50–52, 52–57^ **(Supplementary** Figures 1B-C**; Supplementary Table 1)**.

Together, these results provide a comprehensive genome-wide perspective on the genes and pathways modulated by ER stress and pro-inflammatory cytokines. Comparative analyses suggest that they elicit largely distinct, complementary transcriptional responses, inducing specific response pathways and repressing islet cell type-specific critical functions in response to stress.

### ER stress induces strong and heterogeneous responses in beta cells

To uncover the cell type-specific effects of ER stress and cytokines on islets, we completed single cell (sc) transcriptome profiling of islets (n=3 donors per condition) treated with thapsigargin or pro-inflammatory cytokines **(Supplementary Table 1)**, yielding 18,945 single cell transcriptomes from stressed or control conditions **(Supplementary** Figure 2A**; Supplementary Table 3).** Unsupervised clustering analyses identified each cell type **(Figure 2A; Supplementary Table 3)**, which we annotated using previously reported marker genes such as *GCG* for alpha cells and *INS* for beta cells **(Supplementary** Figures 2A-B**; Supplementary Table 3)**. As expected, alpha (∼38%) and beta cells (∼37%) constituted the majority of islet cells **(Supplementary** Figure 2C**; Supplementary Table 3)**. In striking contrast to the alpha and other islet cell types, beta cells exhibited distinct and increased sensitivity to ER and cytokine stressors, strongly suggested by the identification of a distinct cluster comprised exclusively of stressed beta cells **(Figure 2A)**.

**Figure 2:**
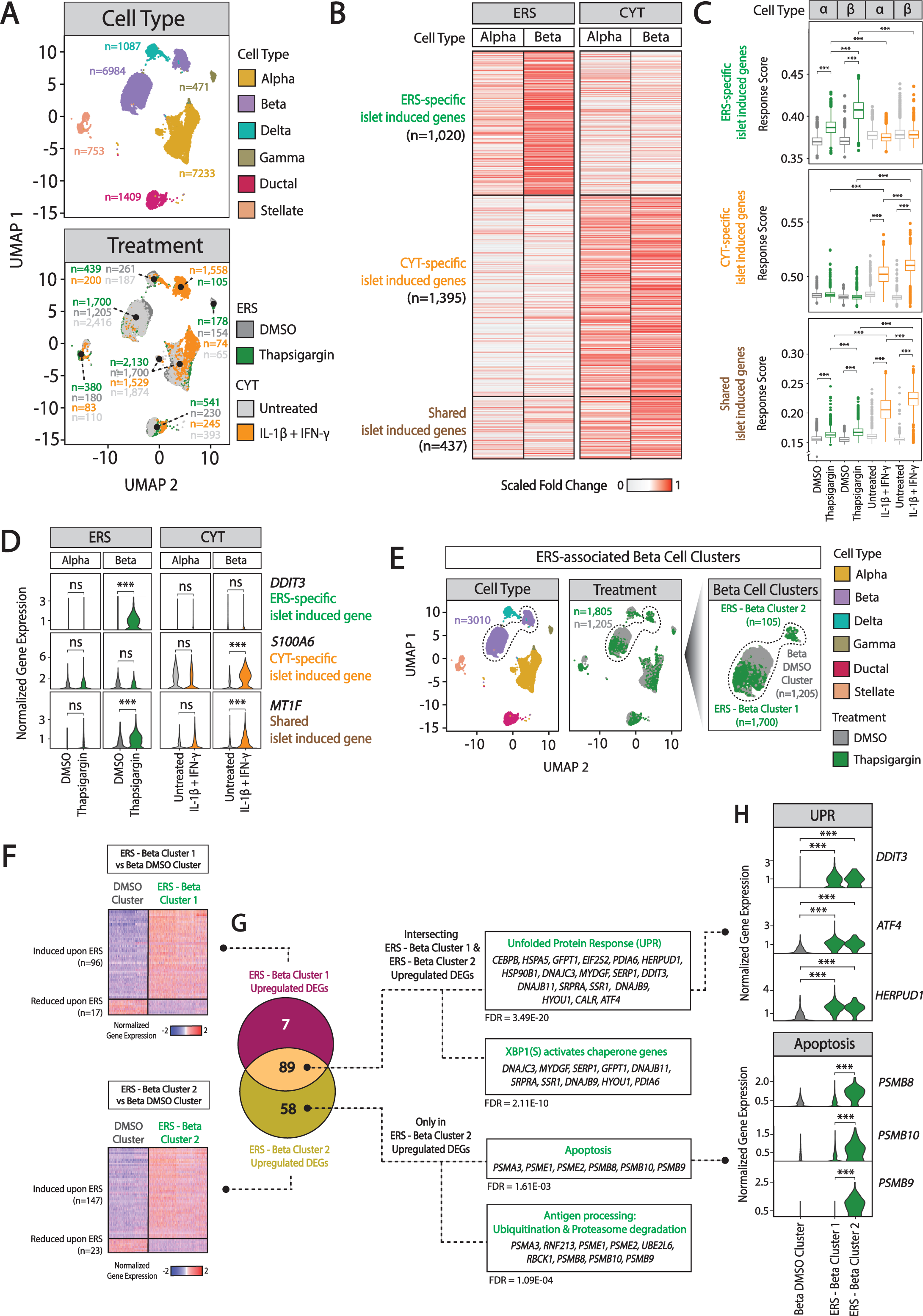
Single-cell transcriptome analysis of human pancreatic islet responses to ER stress (ERS) and pro-inflammatory cytokines (CYT). (A) Uniform Manifold Approximation and Projections (UMAPs) of aggregated single cell transcriptomes from islets exposed to ERS (thapsigargin), CYT (IL-1β + IFNγ) or respective control conditions for 24 hours (n=3 donors per condition; **Supplementary Table 1**). UMAPs are color-coded based on cell type annotations (top) or condition (bottom). n=number of cells per cell type (top) or condition (bottom). **(B)** Scaled fold-change of alpha or beta cell expression of genes induced by ERS or CYT in whole islets. Genes are grouped into genes whose induction is ERS-specific, CYT-specific, or shared between conditions. **(C)** Response scores for the islet-induced genes in alpha and beta cells. ***=p<1.0E-10; ns=not significant, two-sided Wilcoxon test. **(D)** Violin plots of alpha or beta cell expression for representative genes from the three induced gene sets in panel B. ***=FDR<5%, FC≥1.5; ns=not significant. **(E)** UMAP visualization of islet scRNA-seq profiles (left) reveals two beta cell clusters (BC) in ER stressed islets (middle), designated ERS-Beta Cluster 1 (ERS-BC1) or ERS-Beta Cluster 2 (ERS-BC2), respectively (right). The number of cells is indicated in parentheses. **(F)** Heatmaps of significantly induced genes in ERS-BC1 (top) or ERS-BC2 (bottom) versus DMSO control (FDR<5%; FC≥1.5). Number of induced genes in each category is indicated in parentheses. Expression values are scaled using z-scores. **(G)** Venn diagram (left) of significantly induced genes in ERS-BC1 or ERS-BC2 (FDR<5%, FC≥1.5) and the significantly associated pathways from KEGG, Reactome, and WikiPathways for the intersecting *vs.* unique gene sets (right). FDR values for enriched pathways are reported beneath each category. Note that genes specifically induced in ERS-BC2 are significantly associated with apoptosis-related pathways. **(H)** Violin plots showing expression levels of selected unfolded protein response (UPR) or apoptosis genes in ERS-BC1, ERS-BC2, and DMSO control conditions. ***=FDR<5%, FC≥1.5. False discovery rates (FDR) are calculated using Benjamini-Hochberg p-value adjustment. FC, fold-change; DEGs, differentially expressed genes; α, alpha; β, beta.

To determine the relative alpha and beta cell contributions to whole islet transcriptional responses, we assessed expression changes for 1,020 ER stress-specific, 1,395 cytokine-specific, and 437 shared islet stress response genes detected in alpha and beta cell scRNA-seq **(Figure 2B; Supplementary Table 3)**. These data suggested that beta cells exhibit stronger responses to ER stress than alpha cells. To quantify this, we generated ‘response scores’ **(Methods)** using the expression levels of the stress-responsive genes. Interestingly, the response scores showed that, although both alpha and beta cells contribute to ER stress and cytokine responses, beta cells were more likely to yield a response to both ER stress (p<1.0E-10; two-sided Wilcoxon test) and cytokines (p<1.0E-10; two-sided Wilcoxon test) compared to alpha cells **(Figure 2C)**. For example, *DDIT3, S100A6,* and *MT1F* were significantly induced in beta cells but not in alpha cells **(Figure 2D; Supplementary Table 3)**. Similarly, reduced genes detected in ER stressed islets were more significantly (p<1.0E-10; two-sided Wilcoxon test) reduced in beta vs. alpha cells **(Supplementary** Figures 2D-E). For example, genes critical for beta cell function such as *MAFB, SCG2,* and *SHISAL2B*^58–65^ were robustly reduced in beta cells **(Supplementary** Figure 2F**; Supplementary Table 3)**.

Further inspection of the islet scRNA-seq profiles revealed two ER-stressed beta cell subpopulations **(Figure 2E)**, comprising ∼94% (ER stress - Beta Cluster 1 (BC1); n=1,700 cells) vs. ∼6% (ER stress - Beta Cluster 2 (BC2); n=105 cells) of the total ER stressed beta cells **(Supplementary Table 3)**. This heterogeneity in beta cell response was specific to the ER stress condition and was not observed upon cytokine-induced inflammation, nor in alpha cells for either stressor **(Figure 2E, Supplementary** Figure 2G). The distinct ER stressed beta cell subclusters were detected in all 3 donors **(Supplementary** Figure 2H**; Supplementary Table 3)**, suggesting that this is a coherent and robust transcriptional state. To further study the distinct ER stress responses of these beta cell clusters, we compared the transcriptional profiles of each subset to the control condition, which revealed 113 response genes (96 induced; 17 reduced) for ER stress-BC1 and 170 response genes (147 induced; 23 reduced) for ER stress-BC2 **(Figure 2F, Supplementary Table 3)**. 89 (∼58%) of the response genes were shared between the two beta cell subclusters and included *bona fide* ER stress and unfolded protein response (UPR) genes such as *DDIT3*, *ATF4*, and *HERPUD1* **(Figures 2G-H)**. Interestingly, genes induced only in ER stress-BC2 (n=58) were enriched in cellular death-related pathways which consisted of genes in the proteasome superfamily that function to degrade misfolded proteins and regulate apoptosis^64^, such as *PSMB8*, *PSMB9*, and *PSMB10* **(Figures 2G-H)**. These signatures of ubiquitination, degradation, and apoptosis were specific to ER stress-BC2 cluster **(Supplementary** Figures 2I-J**, Supplementary Table 3)**.

In summary, scRNA-seq data revealed that beta cells respond more strongly to ER stress compared to alpha cells. Beta cell responses are composed of two distinct transcriptional states including a smaller subset of beta cells that highly express apoptosis-related genes upon ER stress treatment. This subset of beta cells may represent a distinct beta cell subpopulation that is more sensitive or vulnerable to ER stress-induced cell death or inherent beta cell heterogeneity in the temporal dynamics of ER stress response.

### Identification of ER and inflammatory stress-responsive islet *cis*-regulatory architecture

To determine the *cis-*regulatory elements (CREs) that mediate ER and cytokine stress responses, we mapped and compared genome-wide CRE accessibility in ER or cytokine stressed islets vs. their respective DMSO or untreated controls **(Methods)** using whole islet assay for transposase-accessible chromatin sequencing (ATAC-seq)^65^ **(Supplementary Table 1)**. ∼14% of CREs (14,968/109,399) were significantly (FDR < 5%) remodeled in response to stress; 7,171 CREs were ER stress-responsive (3,375 opening; 3,796 closing) and 8,819 CREs were cytokine-responsive (5,768 opening; 3,051 closing) **(Supplementary Table 4)**. The majority of the responsive CREs exhibited stress-specific accessibility changes **(Figure 3A; Supplementary** Figure 3A**; Supplementary Table 4)**. Among the opening CREs, 2,982 were ER stress-specific, 5,375 cytokine-specific, and only 393 were shared between the two stress conditions.

**Figure 3:**
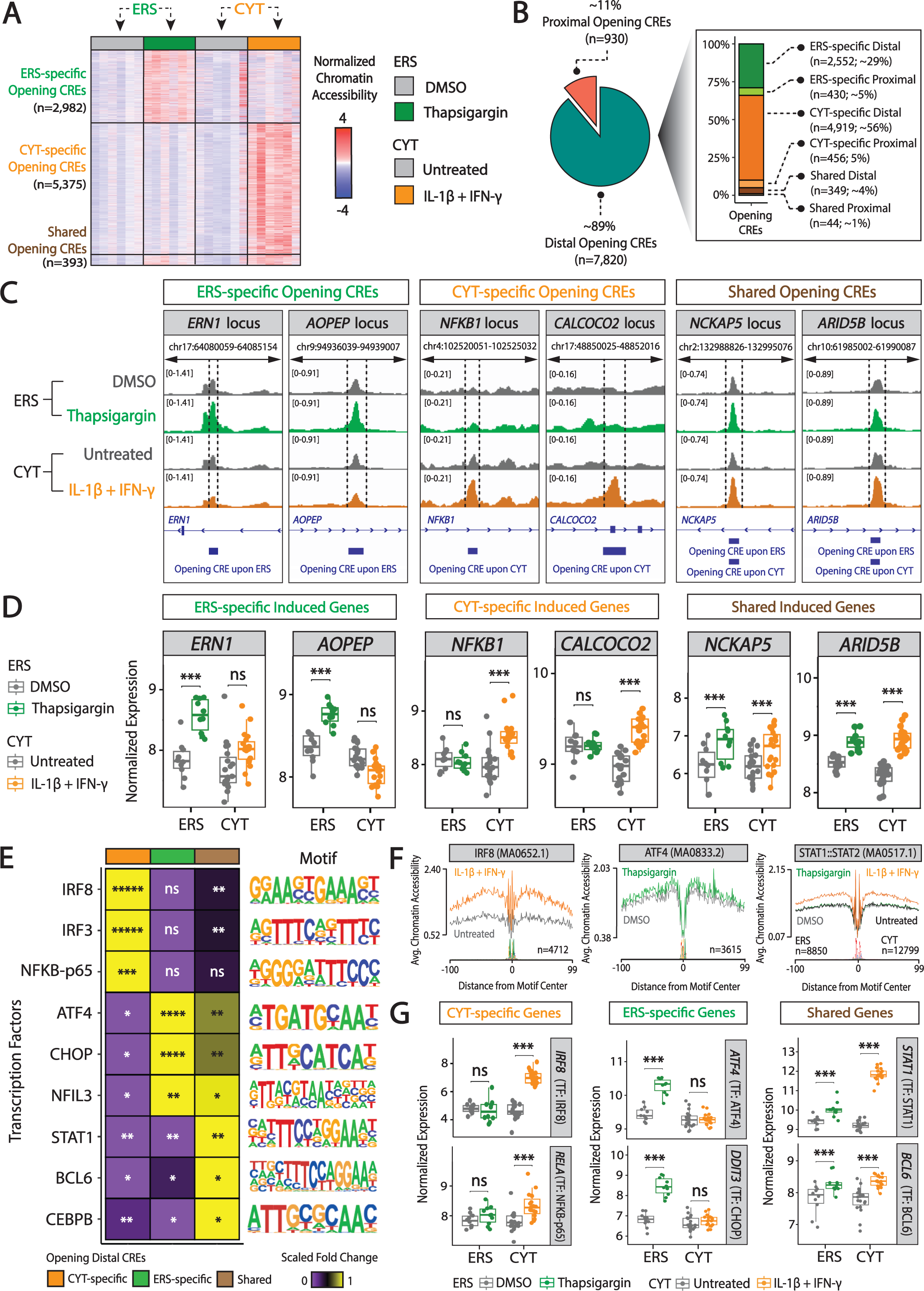
Increased chromatin accessibility changes and associated induced transcriptional regulatory effects of human islet ER stress (ERS) and pro-inflammatory cytokine (CYT) responses. **(A)** Heatmap of human islet *cis*-regulatory elements (CREs) whose accessibility is increased by ERS and/or CYT treatment (FDR<5%). n=number of CREs in each category. Accessibility values are scaled using z-scores. **(B)** Pie chart showing the percent of opening CREs that are proximal vs. distal (≤1kb vs. >1kb to nearest transcription start site (TSS), respectively). Bar chart (right) indicates the percentage of opening CREs per condition. **(C)** Integrative Genomics Viewer (IGV) browser tracks of representative loci highlighting CREs whose increased accessibility is ERS-specific, CYT-specific, or shared. **(D)** Dot-and- box plots of gene expression levels (CPM) per islet donor in treated versus control conditions for responsive genes in representative loci in panel C. ***=FDR<5%, FC≥1.5; ns=not significant. **(E)** Heatmap of enriched transcription factor (TF) motifs identified in ERS-specific, CYT-specific, or shared opening distal CREs. The color gradient indicates the scaled fold change of the motif (i.e., motif instances found in target sequences compared to the background sequences). *****=FDR<1.0E-200; ****=FDR<1.0E-100; ***=FDR<1.0E-50; **=FDR<1.0E-10; *=FDR<1.0E-1; ns=not significant. **(F)** Chromatin footprint analyses indicating average islet chromatin accessibility in IRF8 (left), ATF4 (middle), or STAT1:STAT2 (right) TF binding sites of CYT-specific (left), ERS-specific (middle), or shared opening CREs (right). The number of footprints is indicated with “n=” at the bottom of each footprint plot. **(G)** Dot-and box plots of islet RNA-seq expression levels (CPM) in ERS, CYT, or control conditions for TF-encoding genes with enriched TF motifs or chromatin footprints in panels E or F, respectively. ***FDR<5%, FC≥1.5; ns, not significant. False discovery rates (FDR) are calculated using Benjamini-Hochberg p-value adjustment. FC, fold-change; CPM, counts per million.

The majority of these stress-responsive CREs were distal, i.e., >1kb from transcription start site (TSS)^66,67^ of the nearest expressed gene **(Figure 3B; Supplementary** Figure 3B**; Supplementary Table 4)**, emphasizing the importance of non-promoter CREs in mediating stress responses. We associated the opening and closing distal CREs with the nearest expressed genes in islets and conducted enrichment analyses **(Supplementary Table 4)**. As anticipated, there was a significant correlation between the stress-responsive induced islet chromatin accessibility and gene expression changes (ER stress-specific: p=4.5E-16; cytokine-specific: p=1E-62; shared: p=5.9E-12; Fisher’s exact test) **(Supplementary Table 4)**. For example, we captured ER stress-specific opening CREs in the introns of *ERN1* and *AOPEP* **(Figure 3C; Supplementary Table 4),** genes that were significantly induced by ER stress **(Figure 3D; Supplementary Table 2)**. *ERN1* encodes IRE1α, a central ER stress sensor that initiates UPR and catalyzes unconventional splicing of the ER stress factor XBP1, while *AOPEP* catalyzes N-terminal peptide and amino acid hydrolysis^68–70^. Cytokine-responsive CREs included those within introns of *NFKB1* and *CALCOCO2* **(Figure 3C; Supplementary Table 4)**, two genes that were induced upon cytokine-induced inflammation **(Figure 3D; Supplementary Table 2)**. *NFKB1* is a central mediator of inflammatory responses including in beta cells^71–73^ *CALCOCO2* encodes a selective autophagy receptor and has been recently identified as a putative T2D effector gene that maintains proper beta cell mitochondrial morphology, insulin granule homeostasis, and insulin content^74,75^. Further, we identified increased chromatin accessibility at the *NCKAP5* and *ARID5B* introns upon both stressors, two genes that were induced by both stressors **(Figure 3C-D; Supplementary Tables 2-3)**.

We identified concordant reductions in CREs and nearest gene expression upon ER stress and exposure to cytokines (ER stress-specific: p=9E-53; cytokine-specific: p=3.9E-09; shared: p=3.5E-08; Fisher’s exact test) **(Supplementary Table 4)**. *RAB27B* and *SLC6A17* play key roles in insulin granule exocytosis and amino acid vesicular trafficking, respectively^76–80^. These genes were reduced and linked with closing CREs upon ER stress **(Supplementary** Figures 3C-D**; Supplementary Table 4)**. Similarly, *IGF1R* and *PCSK1*, which play pivotal roles in glucose homeostasis, and proinsulin to insulin processing, respectively^81–83^, were reduced and linked with chromatin closing upon cytokine-induced inflammation **(Supplementary** Figures 3C-D**; Supplementary Table 4)**. *SORL1,* involved in insulin receptor sorting^84^ and *PAX4*, which is crucial for islet development^85,86^ were reduced and linked to chromatin closing upon both stressors **(Supplementary** Figures 3C-D**; Supplementary Table 4)**.

To elucidate the regulatory drivers of islet ER and cytokine stress responses, we identified transcription factor (TF) binding motifs enriched in differential distal peaks **(Supplementary Table 4)**. Motifs for ATF4, CHOP, and NFIL3, which are key transcriptional mediators of UPR^87–90^, were enriched in ER stress-specific opening distal peaks **(Figure 3E).** In contrast, cytokine-specific opening distal peaks were enriched in motifs for interferon response factors IRF8 and IRF3, as well as the NFKB family member NFKB-p65 (alias RELA) **(Figure 3E)**. TF motifs for STAT1, BCL6, and CEBPB were enriched in distal peaks opening upon both stress conditions **(Figure 3E)**. TF motif enrichment analysis for closing distal CREs **(Supplementary Table 4)**, revealed that EOMES, PDX1, and MAFA were enriched in the cytokine-specific, ER stress-specific and shared closing distal CREs, respectively **(Supplementary** Figure 3E). We also observed a concordant downregulation of *EOMES, PDX1,* and *MAFA* under these stress conditions **(Supplementary** Figure 3F). EOMES, PDX1, and MAFA are TFs involved in development- related processes^45,91,92^. The downregulation of these genes, therefore, suggests a potentially coordinated response to stress that could impair the function of the islets, thereby having a significant impact on glucose homeostasis, which can contribute to T2D.

TF footprinting analyses that integrate TF binding motifs with the chromatin accessibility maps^93^ confirmed that, genome-wide, there was a significant increase in chromatin accessibility at the binding sites of ATF4 upon ER stress (p=2.60E-02) and IRF8 upon cytokine-induced inflammation (p=2.95E-04) **(Figure 3F)**. Increased accessibility at the binding sites of these TFs was concordant with the expression changes for the genes encoding these TFs. *ATF4*, *DDIT3,* and *NFIL3* were induced upon ER stress, whereas *IRF8, IRF3* and *RELA* induced upon cytokine-induced inflammation, and *STAT1*, *BCL6*, and *CEBPB* induced by both stressors **(Figure 3G)**.

Together, our ATAC-seq-based analyses reveal that: i) ER stress and cytokine responses in islets substantially remodel the islet epigenome, particularly modulating distal non-coding CREs, and ii) each stressor elicits a distinct epigenetic profile, mediated by different TFs (e.g., CHOP and ATF4 in ER stress; IRF8, NFkB-p65 in cytokines) whose own expression (e.g., CHOP-encoding *DDIT4, ATF4* upon ER stress;) is itself modulated by that stressor.

### T2D-associated genetic variants overlap stress-responsive *cis*-regulatory elements

After comparing ER and cytokine stress-responsive *cis-*regulatory networks, we sought to understand if genetic variants associated with diabetes (T2D/type 1 diabetes (T1D) GWAS) or related glycemic traits might modulate the CREs and processes. Using a set of index and proxy variants **(Methods)** collected from multiple genome-wide studies and meta-analyses^94–102^ **(Supplementary Table 5)**, we identified 212 T2D, T1D, or related glycemic trait-associated variants that overlap stress responsive (opening or closing) CREs **(Figure 4A; Supplementary** Figure 4A**; Supplementary Table 5)**. Twenty-one and 24 T2D- associated variants overlapped ER stress- or cytokine-specific opening CREs, respectively **(Figure 4A; Supplementary Table 5)**. Among these, 11 variants overlapped ER stress-specific opening CREs that are within 500kb of an ER stress response gene **(Figure 4B; Supplementary Table 5),** including *AOPEP* - a key gene involved in peptide processing^70^ and robustly induced by ER stress in beta cells **(Figure 4C; Supplementary Tables 2-3)**. We detected an ER stress-specific induced CRE in the *AOPEP* intron, which harbors the T2D-associated variant rs4744423 **(Figure 4D; Supplementary Tables 4-5)**. The chromatin accessibility of this CRE increased with the T2D risk allele (plus strand: T) of this variant **(Figures 4D-E).** The risk allele is predicted to increase the binding affinity of BATF **(Figure 4F; Supplementary Table 5)**, which is itself an ER stress-responsive islet gene **(Figure 4G; Supplementary Table 2)**. Together, these data suggest that the T2D risk allele rs4744423 is associated with stronger binding of BATF in ER stressed islets, which might lead to increased upregulation of the putative effector gene *AOPEP*. This is supported by the increased expression (p<1.0E-02; two-sided Wilcoxon test) of *AOPEP* in the beta cells of diabetic (T2D) donors compared to non-diabetic (ND) donors **(Figure 4H)** using targeted analysis of human islet single cell transcriptome data we generated in a parallel study^103^.

**Figure 4:**
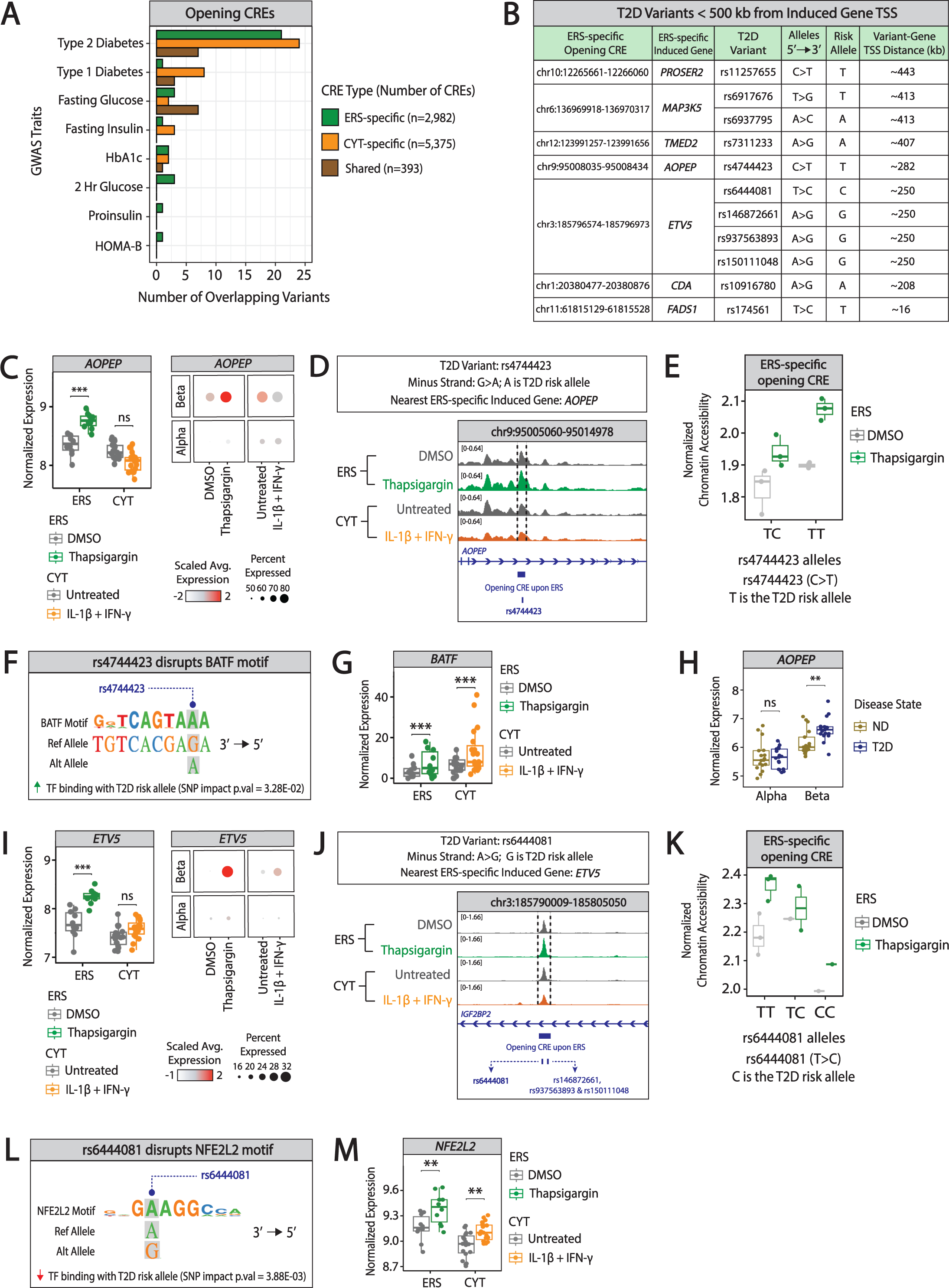
Type 2 Diabetes (T2D)-associated variants overlapping stress-responsive opening CREs. **(A)** Bar chart displaying the number of T2D- or glycemic trait-associated genome-wide association study (GWAS) variants that overlap opening *cis*-regulatory elements (CREs). **(B)** T2D-associated variants overlapping ER stress (ERS)-specific opening CREs located <500 kb from the TSS of an ERS-specific induced gene. **(C)** Expression of *AOPEP*, the putative effector gene of T2D variant rs4744423, under ERS and pro-inflammatory cytokine (CYT) conditions in human islet RNA-seq (left) or scRNA-seq (right) profiles. Dot-and-box plots show gene expression levels (CPM) per islet donor in treated versus control samples. ***=FDR<5%, FC≥1.5 or ns=not significant. Dot plot of *AOPEP* expression in alpha vs. beta cell scRNA-seq profiles in ERS or CYT treated human islets (right). Dot size indicates the percent of *AOPEP*- expressing cells in each cell type; dot color denotes the scaled average *AOPEP* expression in those cells. **(D)** Integrative Genomics Viewer (IGV) browser track showing an ERS-specific opening CRE containing T2D-associated variant rs4744423. **(E)** Dot-and-box plots of islet chromatin accessibility levels (CPM) in donors with rs4744423 TC or TT genotypes (on the plus strand). Note that the homozygous T2D risk allele (TT) genotype is associated with the highest *in vivo* chromatin accessibility. **(F)** Composite logo plot (generated using atSNP^164^) illustrates that the rs4744423 T2D risk allele (T on plus strand, A on minus strand) significantly alters a BATF (indicated by the position weight matrix) transcription factor (TF) binding motif (atSNP p-value=3.33E-02) to create a binding site. **(G)** Expression of *BATF*, the gene encoding BATF, in ERS, CYT or control conditions. Dot-and-box plots show gene expression levels (CPM) per islet donor in treated versus control samples. ***=FDR<5%; FC≥1.5. **(H)** Expression of *AOPEP*, the putative effector gene of T2D-associated variant rs4744423, in human islet alpha and beta cells. Dot-and-box plots show pseudobulk gene expression levels per (CPM) islet donor in cells obtained from non-diabetic (ND) or diabetic (T2D) donors. **=p<1.0E-02; ns=not significant, two-sided Wilcoxon test. **(I)** Expression of *ETV5*, the putative effector gene of T2D-associated variant rs6444081, in ERS, CYT, or control conditions (left). Dot-and-box plots show gene expression levels (CPM) per islet donor in treated vs. control samples. ***=FDR<5%, FC≥1.5; ns=not significant. Dot plot of scRNA-seq data illustrating alpha vs. beta cell *ETV5* expression in ERS or CYT treated human islets (right). Dot size indicates the percent of cells expressing *ETV5* in each cell type; dot color denotes the scaled average expression level of *ETV5* in those cells. **(J)** IGV browser track showing an ERS-specific opening CRE containing T2D-associated variants rs6444081, rs146872661, rs937563893, and rs150111048. **(K)** Dot-and-box plots display islet chromatin accessibility levels (CPM) in donors with rs6444081 TT, TC or CC genotypes (on the plus strand). Note that the homozygous T2D risk allele (CC) genotype is associated with the lowest *in vivo* chromatin accessibility. **(L)** Composite logo plot (generated using atSNP^164^) illustrates that the rs6444081 T2D risk allele (C on plus strand, G on minus strand) significantly disrupts the NFE2L2 (indicated by the position weight matrix) TF binding site (atSNP p-value=3.88E-03). **(M)** Expression of *NFE2L2*, the gene encoding NFE2L2, in ERS, CYT or control conditions. Dot-and-box plots show gene expression levels (CPM) per islet donor in treated versus control samples. **=FDR<5%; FC>1. False discovery rates (FDR) are calculated using Benjamini-Hochberg p-value adjustment. FC, fold-change ; CPM, counts per million.

Similarly, we identified a CRE that was more accessible upon ER stress and harbors the T2D variant rs6444081. The putative effector gene of this CRE is *ETV5*, a modulator of insulin secretion^104–106^, which was induced by ER stress in beta cells **(Figures 4I-J; Supplementary Tables 2-5)**. The T2D risk allele of rs6444081 (plus strand: C) was associated with reduced CRE accessibility **(Figures 4K)** and is predicted to disrupt an NRF2 (encoded by *NFE2L2*) TF binding motif **(Figure 4L; Supplementary Table 5)**, which we previously identified as a putative regulator of islet chromatin accessibility^95^. *NRF2* was induced by ER stress **(Figure 4M)** and, together with KEAP1, it facilitates stress-responsive *ETV5* activation. These data suggest that upon ER stress, the CRE harboring rs6444081 becomes more accessible and regulates *ETV5* activation. However, our data indicate the T2D risk allele rs6444081-C leads to diminished chromatin accessibility, presumably by disrupting NRF2 binding, which would contribute to diminished *ETV5* responses. In alignment, *Etv5^-/-^* mice exhibit impaired insulin secretion and glucose tolerance defects. Knockout islets are smaller and contain smaller beta cells than those from wildtype littermates^104^, and reduced *ETV5* expression was previously reported in T2D vs. ND islets^105^.

Fourteen T2D-associated variants overlapped 11 cytokine-specific opening CREs that are within 500kb of a cytokine-induced gene **(Supplementary** Figure 4B**; Supplementary Table 5),** including *GALNT15* **(Supplementary** Figure 4C**; Supplementary Tables 2-3)** - a member of the GALNT family involved in protein metabolism^107,108^. We detected a cytokine-induced CRE in the intron of *ANKRD28* that harbors the T2D variant rs4685264; the T2D risk allele of rs4685264 (plus strand: G) was associated with increased chromatin accessibility at this CRE **(Supplementary** Figures 4D-E) and increased the binding affinity of the MAX TF **(Supplementary** Figure 4F**; Supplementary Table 5)**, which was induced by cytokines **(Supplementary** Figure 4G**; Supplementary Table 2)**. These data suggest that, upon exposure to cytokines, the CRE harboring rs4685264 becomes more accessible, which allows for increased MAX binding. The T2D risk allele for this variant strengthens predicted MAX binding, potentially leading to the increased upregulation of the putative effector gene *GALNT15*, and ultimately, affecting protein metabolism in islets upon exposure to cytokines. These analyses revealed novel functional roles for T2D variants in modulating cellular responses to ER stress and cytokine-induced inflammation.

### Variant-to-function dissection of ER stress-responsive T2D variant in the *SLC35D3* locus

Integrated analysis of islet multi-omic data from this and previous studies converged to provide new variant-to-function insights for the T2D-associated variant rs6917676, which overlapped an ER stress-responsive, opening CRE that resides in a human islet enhancer hub. This CRE was previously linked to the promoters of nearby genes *MAP3K5*, *SLC35D3*, and *IL20RA*^109^ by promoter capture Hi-C data **(Figure 5A)**. Among these linked genes and other genes in this locus (*MAP7*, *PEX7*, and *IFNGR1*), only *MAP3K5* expression was induced by ER stress, and specifically in beta cells, thereby nominating *MAP3K5* as the likely effector gene of this variant **(Figures 5B-C; Supplementary** Figure 5A**; Supplementary Tables 2- 3)**. The T2D risk allele for rs6917676 (plus strand: T) was associated with increased chromatin accessibility at this ER stress-responsive CRE **(Figure 5D)**. We previously demonstrated that rs6917676 is the expression-modulating variant (emVar) in this CRE using massively parallel reporter assays (MPRA) in mouse MIN6 beta cells, with the T risk allele increasing MPRA activity^102^. To test if the rs6917676-T risk allele is differentially bound by beta cell nuclear/transcription factor(s), we completed electrophoretic mobility shift assays (EMSAs)^110^ using human EndoC-βH3 nuclear extracts **(Figure 5E; Supplementary Table 6)**. EMSA revealed robust T allele-specific binding (red arrows) in untreated, ER stressed, or DMSO solvent control β-cell extracts. The rs6917676-T risk allele is predicted to strengthen an NFIL3 binding motif, **(Figure 5F; Supplementary Table 5),** and the *NFIL3* gene was induced by ER stress in beta cells **(Figure 5G; Supplementary Tables 2-3)**. These data suggest that the T2D risk allele rs6917676-T contributes to islet dysfunction or death by increasing ER stress-responsive *MAP3K5* expression *via* increased NFIL3 binding activity at this ER stress-responsive opening CRE. In alignment, we detected an increased *MAP3K5* expression (p<1.0E-02; two-sided Wilcoxon test) in the beta cells of T2D vs. non-diabetic individuals **(Figure 5H)** using the human islet single cell transcriptome data we analyzed in a parallel study^103^.

**Figure 5:**
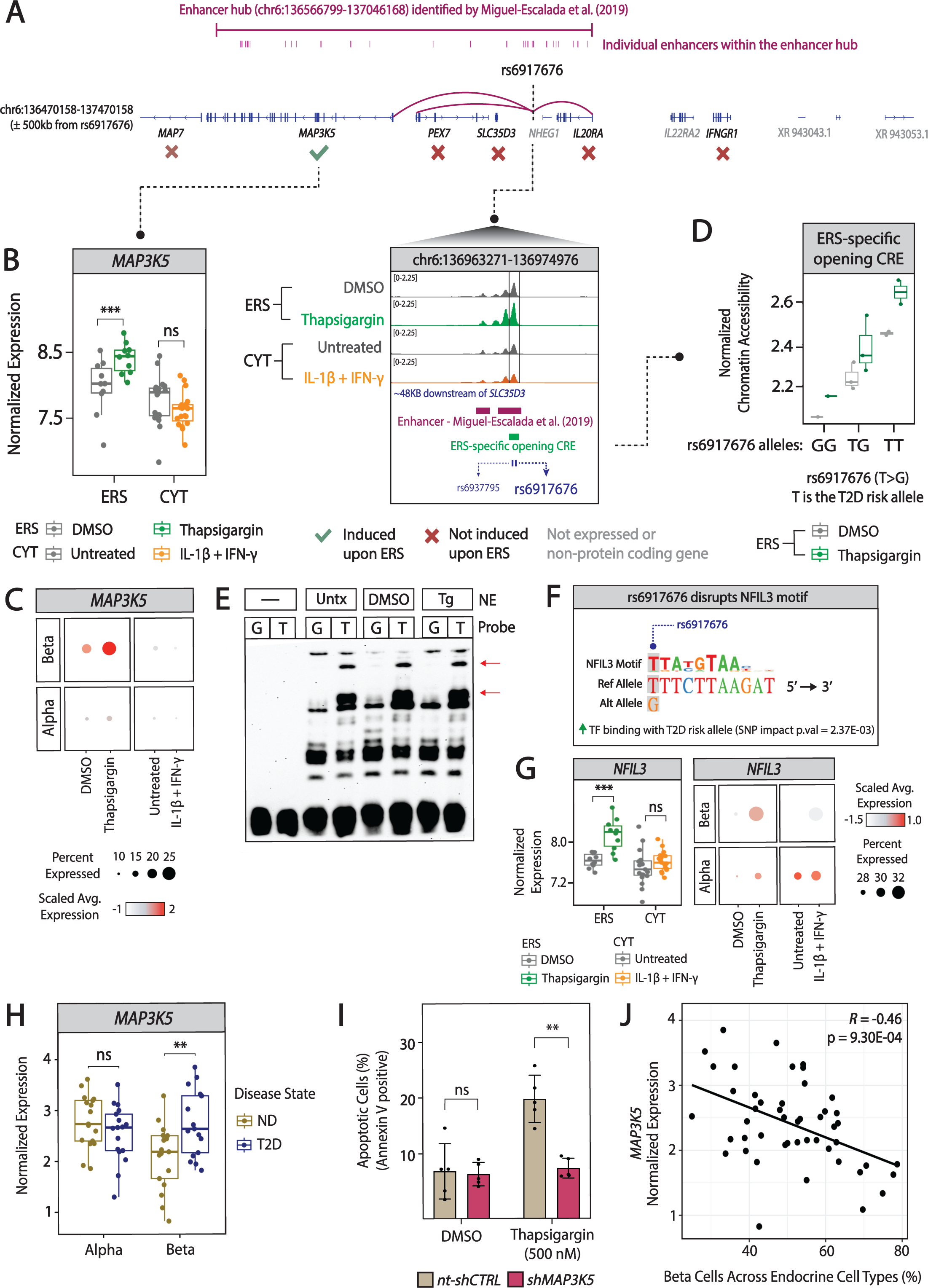
Type 2 Diabetes (T2D)-associated variant rs6917676 potentially modulates beta cell apoptosis in response to ER stress (ERS) via its effector gene, *MAP3K5*. (A) Integrated Genomics Viewer (IGV) browser track showing a ±500kb window (blue gene annotations) centered on T2D- associated variant rs6917676 and an enhancer hub (magenta) identified by Miguel-Escalada et al. (2019)^109^. The enhancer hub encompasses several individual enhancers, with one enhancer (inset) mapping to the ERS-specific, opening distal (≤1kb distance to nearest transcription start site (TSS)) *cis*- regulatory element (CRE) containing T2D-associated variants rs6937795 and rs6917676. All genes located within this ±500kb window are shown; ERS-induced genes are denoted with a green check mark; non-expressed/non-protein coding (grey text) or non-responsive genes are marked by a red “X”. **(B)** Expression of *MAP3K5*, the putative effector gene of T2D-associated variant rs6917676, in human islets in ERS, pro-inflammatory cytokine (CYT), or control conditions. Dot-and-box plots show gene expression levels (CPM) per islet donor in treated vs. control samples. ***=FDR<5%, FC≥1.5; ns=not significant. **(C)** Dot plot of alpha or beta cell MAP3K5 scRNA-seq expression in ERS or CYT treated human islets. Dot size indicates the percent of *MAP3K5* expressing cells in each cell type; dot color represents the scaled average expression level of *MAP3K5* in the cells. **(D)** Dot-and-box plots display islet chromatin accessibility levels (CPM) in donors, stratified by rs6917676 plus strand genotype (GG, TG or TT). Note that *in vivo* chromatin accessibility increases with T2D risk allele (T). **(E)** Electrophoretic mobility shift assay (EMSA) using nuclear extracts (NE) prepared from untreated, DMSO solvent control, or thapsigargin-treated human EndoC-βH3 cells. Red arrows highlight nuclear factors specifically binding the T2D risk allele rs6917676-T. Representative image shown from n=3 EMSAs. **(F)** Composite logo plot (generated using atSNP^164^) illustrates that the rs6917676 T2D risk allele (T on plus strand) significantly alters a NFIL3 (indicated by the position weight matrix) transcription factor (TF) binding motif (atSNP p- value=2.37E-03) to create a binding site. **(G)** Expression of *NFIL3*, the putative effector gene of T2D variant rs6917676, under ERS and pro-inflammatory cytokine (CYT) conditions in human islet RNA-seq (left) or scRNA-seq (right) profiles. Dot-and-box plots show gene expression levels (CPM) per islet donor in treated versus control samples. ***=FDR<5%, FC≥1.5; ns=not significant. Dot plot of *NFIL3* expression in alpha vs. beta cell scRNA-seq profiles in ERS or CYT treated human islets (right). Dot size indicates the percent of *NFIL3*-expressing cells in each cell type; dot color denotes the scaled average *NFIL3* expression in those cells. **(H)** Expression of *MAP3K5*, the putative effector gene of T2D-associated variant rs6917676, in alpha and beta cells. Dot-and-box plots show pseudobulked gene expression levels (CPM) per islet donor in cells obtained from non-diabetic (ND) and diabetic (T2D) donors. **=p<1.0E-02; ns=not significant, two-sided Wilcoxon test. **(I)** Bar plots showing percent of apoptotic (Annexin V-positive) cells detected in human EndoC-βH3 cells exposed to 500nM thapsigargin or DMSO solvent control (Annexin V staining) after *MAP3K5* knockdown (*shMAP3K5*) vs. non-targeting *shRNA* control (*nt-shCTRL*). n=5 biological replicates per condition. **=p<1.0E-02; ns=not significant, two-tailed t-test. **(J)** Plot of the correlation between normalized *MAP3K5* expression (CPM) and the proportion of endocrine cells that are beta cells for 48 human islet donors (Motakis and Nargund et al., *in preparation*^103^). Note the statistically significant inverse relationship between *MAP3K5* expression and beta/endocrine percentages. False discovery rates (FDR) are calculated using Benjamini-Hochberg p-value adjustment. FC, fold-change; Untx, untreated; Tg, thapsigargin.

*MAP3K5* encodes the MAPK kinase ASK1, which activates JNK and p38 in stress responses^111^. ASK1 is activated by ER stress in MIN6 β cells, and *Ask1/Map3k5* knockdown or germline deletion increases MIN6 cell survival and reduces islet caspase activity, respectively^112^. To test if *MAP3K5* modulates ER stress-responsive apoptosis in human beta cells, we assessed how *MAP3K5* shRNA knockdown altered apoptosis in EndoC-βH3 cells exposed to a (patho)physiologic range of thapsigargin concentrations (125- 2000 nM). We achieved approximately 80% knockdown of *MAP3K5* (**Supplementary** Figure 5B). *MAP3K5* deficient cells exhibited significantly fewer apoptotic (Annexin V-positive) cells compared to the non-targeting shRNA control cells exposed to pathophysiologic thapsigargin concentrations **(Figure 5I; Supplementary** Figure 5C**; Supplementary Table 6)**. Interestingly and consistent with its pro-apoptotic role in ER-stressed EndoC-βH3, we found that increased *MAP3K5* expression was significantly associated with reduced beta cell in human islets **(Figure 5J).** Taken together, these data suggest that *MAP3K5* plays a pivotal role in modulating ER stress-induced beta cell apoptosis and that the T2D- associated rs6917676-T risk allele contributes to T2D risk or progression by enhancing ER stress-responsive *MAP3K5* expression.

## DISCUSSION

This study provides novel genome-wide insights into the transcriptional regulatory circuitry mediating pancreatic islet stress responses, particularly to ER stress and pro-inflammatory cytokines, two pathophysiologic stressors implicated in T2D pathogenesis. Through comprehensive RNA-seq and ATAC-seq analyses, we identified distinct sets of genes and CREs that are responsive to ER stress and cytokine-induced inflammation. The majority of stress-responsive genes and CREs were specific to either ER stress or cytokines. Using scRNA-seq, we uncovered alpha and beta cell specificity of these responses. The context-specific responses of islets to ER stress and cytokines are intriguing yet not entirely unexpected. The specificity likely reflects a finely tuned cellular mechanism, which allows for islets to adapt to and tailor their responses to diverse pathophysiologic stimuli. For example, we found that ER stress predominantly triggered pathways related to protein folding and secretion, which are crucial for beta cells’ insulin- producing function^113^. In contrast, cytokine treatment activated pro-inflammatory and signaling pathways that can interfere with crucial islet function such as insulin secretion^36,114^.

scRNA-seq profiling of these stress responses in human islets revealed cell type-specificity of responses to ER stress and cytokine-induced inflammation. Beta cells responded more substantially than alpha cells to both stressors. These data also uncovered heterogeneity in beta cell responses to ER stress, marked by the presence of two transcriptionally distinct heterogeneous beta cell subpopulations. One of these subsets (ER stress-BC1) reflected the activation of *bona fide* ER stress response genes and pathways (e.g., *DDIT3* and *ATF4*) whereas the other smaller subset (ER stress-BC2) also included the induction of apoptosis-related genes (e.g., *PSMB8* and *PSMB9*). Interestingly, this apoptotic beta cell subpopulation was detected in all donors. These findings suggest that a fraction of beta cells are prone to ER stress-induced cell death, which could contribute to beta cell death associated with T2D^115,116^.

Epigenetic responses to these stressors mostly occurred in the distal regulatory regions of the genome, highlighting the importance of the noncoding genome in cellular responses to stress. Stress-responsive opening CREs were enriched in binding sites for critical TFs (e.g., ATF4 upon ER stress, IRF8 upon cytokine-induced inflammation). Genes encoding these TFs were also activated upon these stressors, suggesting that cellular responses are tightly regulated at the epigenetic level by the activation of critical TFs as well as by the increased chromatin accessibility at their binding sites. By intersecting T2D- associated genetic variants with stress-responsive CREs, we uncovered 52 variants residing in 38 ER or cytokine stress-induced CREs, suggesting that these candidate functional T2D variants contribute to T2D etiology by altering these responses. Although the identification of stress-responsive CREs, their overlap with T2D-associated variants, and targeted allelic analyses implicate this subset of T2D variants as genetic modulators of these responses, larger sample sizes are needed to formally demonstrate their allelic effects on stress-responsive chromatin accessibility and gene expression using allelic imbalance or quantitative trait locus approaches. Additionally, the exploration of additional T2D-associated pathophysiologic stressors (e.g., glucolipotoxicity or oxidative stress) or various stimuli, and their interaction with genetic variants, could further help elucidate the complex molecular landscape of T2D, stratify T2D association variants/signals into functional bins, and identify new therapeutic gene targets and pathways.

Taken together, our data and analyses uncovered novel functional associations of T2D variants in modulating cellular responses to ER stress and cytokine-induced inflammation. Among these variants, we identified an ER stress-responsive CRE that contains the T2D variant rs6917676. Human islet pcHi-C data from Ferrer and colleagues and RNA-seq from this study converge to nominate *MAP3K5* as the target gene of this CRE and the T2D effector gene for this genetic association signal. *MAP3K5* encodes MAP3K5 (alias ASK1), a kinase that promotes apoptosis *via* activation of JNK and p38 signaling pathways^112,117–119^. This association suggests a potential mechanism in which the rs6917676 T2D risk allele enhances ER stress-induced beta cell *MAP3K5* expression, which promotes excessive apoptosis to exacerbate beta cell loss in T2D. In alignment with this, *MAP3K5* expression levels were inversely correlated with beta cell abundance in a 48-donor islet scRNA-seq cohort, and T2D donors in this cohort had a higher *MAP3K5* expression and significantly fewer beta cells comprising their islets^103^. Selonsertib, a MAP3K5 (ASK1) inhibitor, and its structural analog GS-444217^120,121^ have been shown to improve diabetic nephropathy by targeting p38 in pre-clinical rodent models of diabetes^122–124^. Randomized placebo-controlled double-blind Phase 2 clinical trials (Clinical Trial Identifier: NCT04026165)^125^ for diabetic complications, such as diabetic kidney disease^126,127^, have been successful, and Selonsertib has now been approved for Phase 3 clinical trials to prevent/treat moderate to advanced diabetic nephropathy^124,128^. Our data suggests that this compound might also be an effective primary intervention to combat progression to T2D by preserving mass and function of ER-stressed beta cells. More broadly, these findings highlight the significance of studying GWAS variants in the context of stress conditions, which more closely reflect cellular state during disease, including for T2D.

In conclusion, this comprehensive and comparative multi-omic mapping study provides important new mechanistic insights into how human islet cells respond to two important stressors: ER stress and cytokine-induced inflammation. Importantly, these maps enabled the nomination of new candidate causal T2D-associated genetic variants that likely contribute to T2D risk or progression by modulating these responses. These findings support the growing literature emphasizing the importance of cell- and context-specific responses in the pathophysiology of and approaches to combat islet dysfunction in T2D. Our study not only enhances our understanding of T2D pathogenesis, but also offers potential new genetics-based avenues or insights, such as repurposing ASK1 inhibitors to combat ER stress-induced beta cell apoptosis, for targeted interventions to preserve beta cell function under pathophysiologic ER stress.

## MATERIALS AND METHODS

### Study Subjects and Primary Islet Culture

Fresh human cadaveric pancreatic islets were procured from Prodo Labs or the Integrated Islet Distribution Program (IIDP) **(Supplementary Table 1)**. Upon arrival, cells were transferred into PIM(S) media (Prodo Labs) supplemented with PIM(ABS) (Prodo Labs) and PIM(G) (Prodo Labs) and incubated in a T-150 non-tissue culture treated flask (VWR) for recovery at 37°C and 5% CO2 overnight. The following day, media was changed to CMRL (10% FBS, 1% Glutamax) supplemented with either 0.025%v DMSO, 250nM thapsigargin or 25 U/mL of IL1β + 1000 U/mL of IFNγ (R&D Systems). After 24-hr incubation at 37°C and 5% CO2, nuclei and total RNA were isolated for RNA-seq and ATAC-seq library preparation as previously described^95^.

### RNA-seq Library Preparation and Sequencing

Human islet RNA-seq libraries were prepared from total RNA using the stranded TruSeq kit (Illumina). ERCC Mix 1 or Mix 2 spike-ins were randomly added to each sample (Thermo Fisher, catalog #4456740) before pooling and sequencing on Illumina NovaSeq S4 to an average depth of 50 million paired-end reads per sample as previously described^95^. The paired-end (2x150 bp) RNA-seq FASTQ files for each islet were aligned against the human genome (GRCh38/hg38) using STAR^105,129^ and counts were generated using QoRTs^130^ **(Supplementary Table 2)**.

### RNA-seq Analyses

Genes were annotated using Ensembl^131^, and only genes in autosomal chromosomes were considered for downstream analysis. Non-protein coding genes in autosomal chromosomes, including RNA and pseudogenes (annotated as ‘transcribed_unprocessed_pseudogene’, ‘processed_pseudogene’, ‘lncRNA’, ‘unprocessed_pseudogene’, ‘TR_V_pseudogene, snRNA’, ‘misc_RNA’, ‘rRNA_pseudogene’, ‘IG_V_pseudogene’, ‘IG_C_pseudogene’, ‘TEC, scRNA’, ‘translated_processed_pseudogene’, ‘vault_RNA’, ‘sRNA’, ‘pseudogene’, ‘transcribed_unitary_pseudogene’, ‘transcribed_processed_pseudogene’, ‘unitary_pseudogene’, ‘miRNA’, ‘snoRNA’, ‘rRNA’, ‘TR_J_pseudogene’, ‘ribozyme’, ‘IG_J_pseudogene’, ‘scaRNA’, ‘translated_unprocessed_pseudogene’, and ‘IG_pseudogene’) were filtered out. The remaining (protein coding) genes were then filtered for expression by requiring >0 CPM in ≥8 samples, and ERCCs were filtered for expression by requiring >5 reads in ≥2 samples. Normalization of protein coding genes with ERCC was performed using RUVSeq^132^, which also estimated unwanted variation (W_1) in the data. Surrogate variable analysis was then performed using svaseq^133^, and the surrogate variables that explained >10% of variance in the data (n=3) were considered in downstream analysis. Genes were then tested for differential expression (FDR<5%; |LFC|≥0.585) between their respective control (DMSO; untreated) and treatment (thapsigargin; IL-1β+IFN-γ) conditions **(Supplementary Table 2)**, with gene expression adjusted for age, sex, batch, BMI, surrogate variables and W_1, using edgeR’s^134^ ‘tagwise’ and robust dispersion estimation parameter on TMM normalized counts. FDR was calculated using Benjamini-Hochberg p-value adjustment. The differentially expressed genes were classified as specific or shared using a Venn diagram, and were input into DAVID^135^ to find the enriched pathways (FDR<10%) using KEGG^136^, Reactome^137^, and WikiPathways^138^ **(Supplementary Table 2)**.

### Single Cell RNA-seq Library Preparation and Sequencing

After a 24-hour treatment, as described above, islets from six organ donors **(Supplementary Table 1)** were treated with Accutase for 8-10 min at 37°C to generate a single cell suspension. Cells were then washed and suspended in Staining buffer (PBA, 2%BSA, 0.01%TweenS), and immediately processed as follows: incubated with Fc Blocking reagent (FcX, BioLegend) for 10 minutes at 4 °C, incubated with 0.5ug of a unique Cell Hashing antibody (TotalseqTM-A0251 to A0257 anti-human hashtag antibody, BioLegend) for 20 minutes at 4 °C, and washed two times with Staining buffer and once with PBS+0.04%BSA. Cell viability was assessed on a Countess II automated cell counter (ThermoFisher), and up to 30,000 cells (∼5,000 cells from each hash-tagged (HTO) sample) were loaded onto one lane of a 10X Chromium Controller. One single-cell suspension was loaded twice, i.e. onto two lanes of a 10x chip. Two gene expression and two HTO libraries were generated per islet sample, which were combined into a single set of gene expression and HTO outputs. Single cell capture, barcoding and library preparation were performed using the 10X Chromium platform V3 chemistry and according to the manufacturer’s protocol (#GC000103). cDNA and libraries were checked for quality on Agilent 4200 Tapestation, quantified by KAPA qPCR, and pooled and sequenced on an Illumina NovaSeq 6000 S2/S4 flow cell lane, targeting an average sequencing depth of 50,000 reads per cell. Illumina base call files for all libraries were converted to FASTQ using Illumina’s bcl2fastq^139^. The FASTQ files were then associated with the gene expression libraries, aligned to the GRCh38.93 reference genome and merged, including all transcribed unitary pseudogenes, using the 10x Genomics Cell Ranger’s count pipeline^140,141^. FASTQ files representing the HTO libraries were processed into hashtag-count matrices using CITE-seq-Count^142^ **(Supplementary Table 3)**.

### Single Cell RNA-seq Clustering and Annotation

Sample identities were determined using demuxlet^143^. Ambient RNA for each islet was removed using SoupX^144^ by setting contamination fraction to 20%. SoupX-adjusted data were then demultiplexed based on enrichment of HTO using Seurat^145^. Only cells with genes >2000 and mitochondrial percentage <40% were considered for downstream analysis. Doublet cells were then identified using Scrublet^146^ and removed. To filter out any remaining potential doublets or multiplets, cells in the >0.95 quantile with respect to the number of genes expressed were removed **(Supplementary Table 3)**. These data were then merged into a single object using Seurat^145^, and corrected for batch effect using Harmony^147^. Seurat’s^145^ ‘FindClusters’ was implemented to identify cell clusters, which were then annotated for cell type identity using islet marker genes **(Supplementary Table 3)**. Seurat clusters that expressed more than one marker gene were classified as doublets and removed from downstream analyses.

### Single Cell RNA-seq Data Analyses

To generate response scores, the differentially expressed genes from bulk data were curated into ER stress-specific, CYT-specific and shared response gene modules. UCell’s^148^ ‘AddModuleScore_UCell’ was then used to calculate each module’s enrichment (i.e. response) scores **(Supplementary Table 3)**. To identify expressed genes between the control (DMSO; untreated) and treatment (thapsigargin; IL- 1β+IFN-γ) conditions in alpha and beta cells, Seurat’s^145^ ‘FindMarkers’ was implemented using the MAST^149^ test and adjusted with respect to batch and disease state **(Supplementary Table 3)**. This methodology was also implemented on only those genes that were detected in a minimum of 10% of cells in either of the beta cell subpopulations to identify differentially expressed genes between ER stress-BC1 vs. DMSO (FDR<5% and |LFC|≥0.585), ER stress-BC2 vs. DMSO (FDR<5% and |LFC|≥0.585), and ER stress-BC1 vs. ER stress-BC2 (FDR<5%) comparisons **(Supplementary Table 3)**. These differentially expressed genes were then input into DAVID^135^ to find the enriched pathways (FDR<10%) using KEGG^136^, Reactome^137^, and WikiPathways^138^ **(Supplementary Table 3)**.

### ATAC-seq Library Preparation and Sequencing

Human islet ATAC-seq libraries were prepared following the Active motif ATAC prep kit (Active motif catalog# 53150). Briefly, 50 islet equivalents (50,000 cells) per sample were transposed in triplicate, libraries were barcoded, pooled into 3-islet batches, and sequenced using 2 x 150 bp Illumina NovaSeq S4 chemistry as previously described^95^. The paired-end (2x150 bp) ATAC-seq FASTQ files for each islet were trimmed using Trimmomatic^150^, and aligned against the human genome (GRCh38/hg38) using BWA- MEM^151^. Duplicate reads were removed, and the remaining reads were shifted as previously described^152,153^. Using SAMtools^154^, technical replicates were merged and peaks were called using MACS2’s^155^ ‘BAMPE’ parameter. TDF files were generated using IGVTools^156^ to visualize peaks on IGV^156^. Separate consensus peaksets for ER stress and CYT samples were generated by considering peaks that were present in at least two samples; peaks mapping to ENCODE Exclusion List Regions^157^ were removed using DiffBind^158^. The union of all peaks from ER stress and CYT samples was determined using GenomicRanges^159^, and counts were normalized using CPM **(Supplementary Table 4)**.

### ATAC-seq Data Analyses

Only peaks in autosomal chromosomes were considered, which were then filtered for depth by requiring >0 CPM in ≥8 samples. Surrogate variable analysis was then performed using svaseq^133^, and the surrogate variables that explained >10% of variance in the data (ER stress: n=2; cytokines: n=3) were considered in downstream analysis. Peaks were then tested for differential accessibility (FDR<5%) between their respective control (DMSO; untreated) and treatment (thapsigargin; IL-1β+IFN-γ) conditions **(Supplementary Table 4)**, with accessibility adjusted for age, sex, batch, BMI, and surrogate variables using the edgeR^134^ ‘tagwise’ and robust dispersion estimation parameter on TMM normalized counts. FDR was calculated using Benjamini-Hochberg p-value adjustment. Peaks were then annotated to the nearest expressed protein-coding gene extracted from GENCODE v35^160^ in islets using HOMER’s^161^ ‘ annotatePeaks.pl’ command. Peaks with distance ≤1kb to the nearest expressed gene’s TSS were considered proximal, and the other peaks were considered distal^66,67^. IRange’s^159^‘subsetByOverlaps’ function was used to classify proximal and distal peaks as specific or shared **(Supplementary Table 4)**.

### Enrichment and Footprinting Analysis

Nearest genes to differentially accessible peaks were used as input into DAVID^135^ to find the enriched pathways (FDR<10%) using KEGG^136^, Reactome^137^, and WikiPathways^138^ **(Supplementary Table 4)**. TF motifs present in the differentially accessible peaks were found using HOMER’s^161^ ‘findMotifsGenome.pl’ command. FDR of TFs was calculated using Benjamini-Hochberg p-value adjustment, and the fold change was calculated by dividing ‘% of Targets Sequences with Motif’ by the ‘% of Background Sequences with Motif’ **(Supplementary Table 4)**. Using SAMtools^154^, samples of the same control or treatment conditions were merged, and peaks were called using MACS2^155^ with ‘BAMPE’ parameter. HINT-ATAC^93^was used to identify TF footprints and to calculate differences in TF activity between the respective control and treatment conditions **(Supplementary Table 4)**.

### Overlapping Genetic Variants with Peaks

Index variants associated with T1D, T2D, and glycemic traits (fasting glucose, fasting insulin, HbA1c, 2- hour glucose, HOMA-B, HOMA-IR, proinsulin, modified Stumvoll insulin sensitivity index, and disposition index) were obtained from the largest and most recent genome-wide association meta-analyses for each trait^94–102^(**Supplementary Table 5**). Proxy variants, or variants in strong linkage disequilibrium (LD) with the index variant, were defined as any variant that was in LD r2≥0.75 with the index variant calculated using the 1000 Genomes Phase 3 reference panel^162^ which was accessed through ‘https://ldlink.nih.gov’ using the global ancestry group that most closely matched the original GWAS meta-analysis; all individuals in the reference panel were used for GWAS meta-analyses of multi-ancestry populations. When necessary, index and proxy variants were lifted over to hg38 genome; variants that we were unable to lift over were not included in our analyses **(Supplementary Table 5)**. Index and proxy variants with a reference SNP ID (rsID) assigned by dbSNP^163^ were then overlapped with differentially accessible peaks using IRange’s^159^ **‘**findOverlapPairs’ function to find T2D variants that are harbored by stress-responsive peaks **(Supplementary Table 5)**. T2D variants that overlapped stress-responsive peaks and were located <500kb from the nearest induced or reduced gene were then entered into atSNP^164^ to identify all TF motifs being disrupted by the variant in the sense or antisense strands, and only those motifs with a ‘SNP impact p-value’ <0.05 were considered downstream. This list of motifs was then cross-referenced against the list of enriched TF motifs identified by HOMER^161^ (as described above) to determine relevant TF motifs **(Supplementary Table 5)**. ATAC-seq read pileups were used to infer the genotypes of donors for the T2D variants using pysam^165,166^ **(Supplementary Table 5)**.

### EndoC-βH3 Cell Culture

EndoC-βH3 cells were cultured in Advanced DMEM F-12 media (Invitrogen) containing 2% BSA (Sigma), 2mM Glutamax (Gibco), 50uM 2-beta mercaptoethanol (Sigma), 10mM nicotinamide (SIGMA), 6.7ng/ml sodium selenite (Sigma), 1% Penicillin/Streptomycin (Gibco) and 10ug/ml Puromycin (Calbiochem) on ECM (Sigma) and Fibronectin (Sigma) coated flasks^167^.

### shRNA knockdown in EndoC-βH3

Plasmid pLKO-puro shRNA clones (Mission shRNA) were purchased from Sigma (SHC016 (shCTRL); TRCN0000000993 (shMAP3K5). Lentivirus was produced in HEK293T cells co-expressing the shRNA plasmid together with psPAX2 packaging plasmid and pVSV-G envelope plasmid (Addgene). Virus was concentrated using Lenti-X Concentrator (Takara) and titer quantified using p24 ELISA antigen assay (Takara). MOI=5 was used to transduce 1x10^97^ EndoC-βH3 cells in culture media without pen/strep and puromycin.

Cells were collected for RNA extraction 96 hrs post transduction using TRIZOL (Invitrogen), phase separation was achieved using Chloroform. Isopropanol was used for RNA precipitation using glycogen as a carrier, the pellets were washed using 75% ethanol, air-dried, and resuspended in DEPC water. RNA was measured using Qubit RNA HS Assay (ThermoFisher). Total RNA was used to perform qPCR using RNA to CT kit (Invitrogen) and FAM-Taqman probes (Invitrogen) and analyzed on QuantStudio 7 (Applied Biosystems) normalized to TBP/HPRT1 Taqman probe **(Supplementary Table 6).**

### Flow cytometry analysis of beta cell apoptosis

Eighteen hours post-transduction, media was changed to pen/strep and puromycin complete media with 0, 125, 250, 500, 1000, 1500, 2000nM thapsigargin (Sigma Aldrich) dissolved in DMSO or 0.5% DMSO solvent control (VWR). Ninety hours after transduction, cells were collected using Trypsin (Gibco) and stained using PE-Annexin V Apoptosis Detection Kit (BioLegend) according to manufacturer’s instructions. The samples were assessed on Fortessa (BD Sciences) and annexin V-positive cells were analyzed and quantified using FlowJo Software (BD Sciences) **(Supplementary Table 6).**

### EMSA

Electrophoretic mobility shift assays (EMSAs) were carried as previously described^110^. Nuclear extracts were prepared from EndoC-βH3 cells using NE-PER Extraction kit (Thermo Fisher Scientific), quantified using Pierce BCA protein assay kit (Thermo Fisher Scientific), and stored in -80°C until use. Twent-one- bp biotin end-labeled, complementary oligonucleotides were designed to the variant rs6917676 (5’-bio- TAATGACTGT[**G/T**]TTCTTAAGAT-3’, Integrated DNA Technologies), and double stranded probes were generated for both alleles. The Lightshift EMSA optimization and control kit (Thermo Fisher Scientific) was used according to the manufacturer’s instructions. Each reaction consisted of a 10x binding buffer, Poly Di-Dc, 4µg of nuclear extract, and 200nM of labeled probe. Reactions were incubated at 25°C for 25 minutes. DNA-protein complexes were detected using Lightshift Chemiluminescent Nucleic Acid Detection kit (Thermo Fisher Scientific) according to manufacturer’s protocol. EMSAs were repeated at least three times and yielded comparable results.

### DATA AVAILABILITY

All cadaveric human islet ATAC-seq, RNA-seq, and single cell RNA-seq data generated in this study are available via the Gene Expression Omnibus under study accession GSE251913.

## ACKNOWLEDGEMENTS

Human pancreatic islets and/or other resources were provided by the NIDDK-funded Integrated Islet Distribution Program (IIDP) (RRID:SCR_014387) at City of Hope, NIH Grant # 2UC4DK098085; we are gratefully indebted to the anonymous islet organ donors and their families. We gratefully acknowledge the contribution of the Single Cell Biology service, the Genome Technologies service, and Research Cyberinfrastructure computational resources at The Jackson Laboratory for expert assistance with the work described in this publication. We thank members of the Ucar and Stitzel labs for critical feedback during the progress of this study, Taneli Helenius for aid in manuscript revisions, and Natalia A. Mozzicato for organizing the references and citations. Graphical abstract was created with BioRender.com. This study was made possible by the generous financial support of the United States Department of Defense (DOD) under award number W81XWH-18-0401 (to MLS, DU) and the National Institutes of Health (NIH) under award number R01DK118011 (to MLS), R01AG052608 (to DU), and U01AI165452 (to DU). CNS was also funded through financial support from the American Diabetes Association 11-22-JDFPM-06. Opinions, interpretations, conclusions, and recommendations are solely the responsibility of the authors and do not necessarily represent the official views of the NIH or DOD.

## AUTHOR CONTRIBUTIONS

D.U. and M.L.S. designed and secured funding for this study. R.K. and R.M.B. coordinated human islet sample collection, preparation, and data generation. E.K.S. analyzed the data. E.K.S., D.U., and M.L.S. interpreted the data and wrote the manuscript. V.S. performed EMSA and shRNA knockdown analyses. C.Z. and C.N.S. created and curated the lists of GWAS T2D, T1D, and glycemic trait variants from previous studies. A.T. helped with computational analyses. All authors read and revised the manuscript, figures, and tables prior to submission.

