## Supplementary figures and images for "Comparative Multi-omic Mapping of Human Pancreatic Islet Endoplasmic Reticulum and Cytokine Stress Responses Provides Insights into Type 2 Diabetes Genetics"

### Supplementary Figure 1

SUPPLEMENTARY FIGURE 1

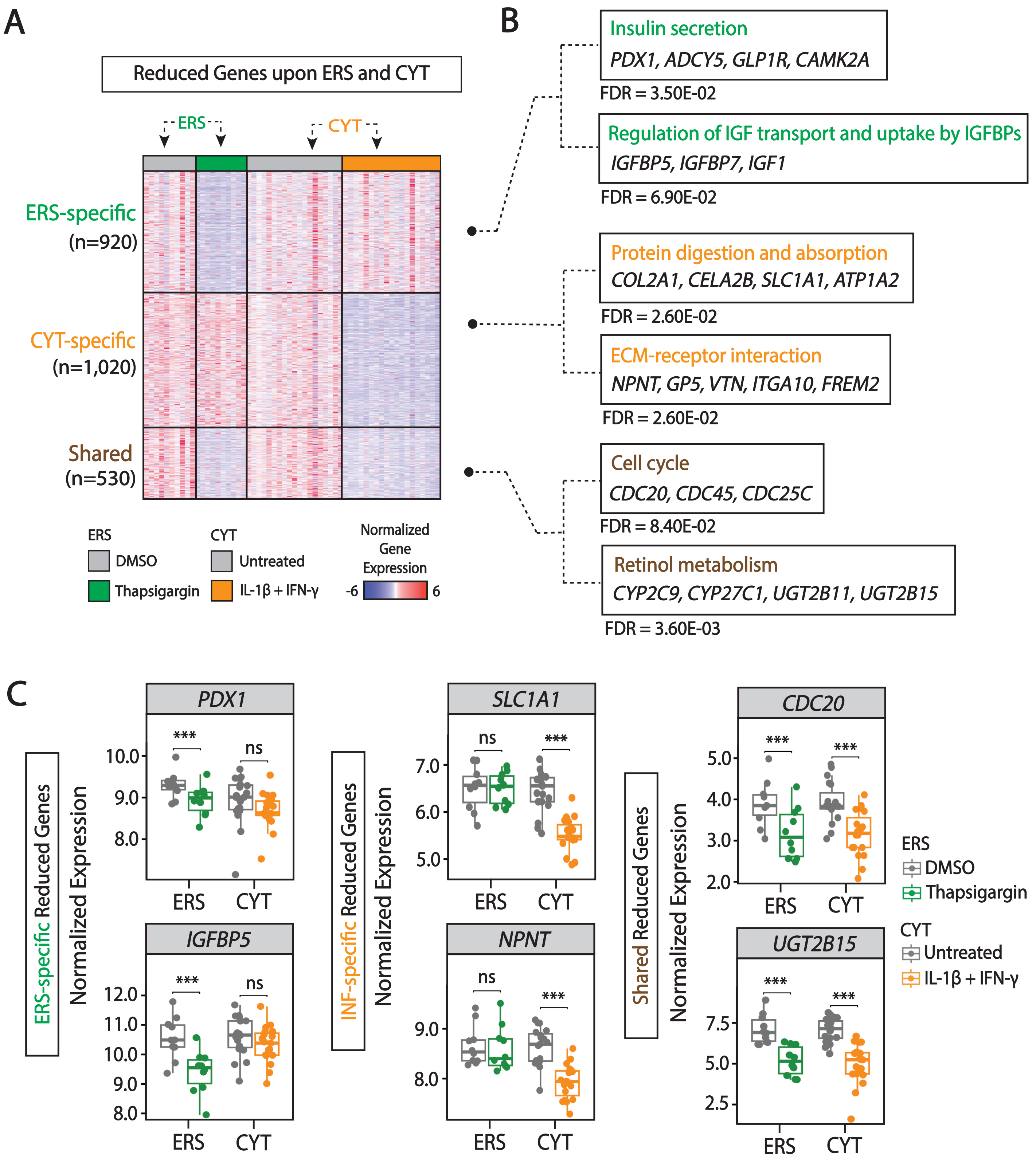

### Supplementary Figure 2

SUPPLEMENTARY FIGURE 2

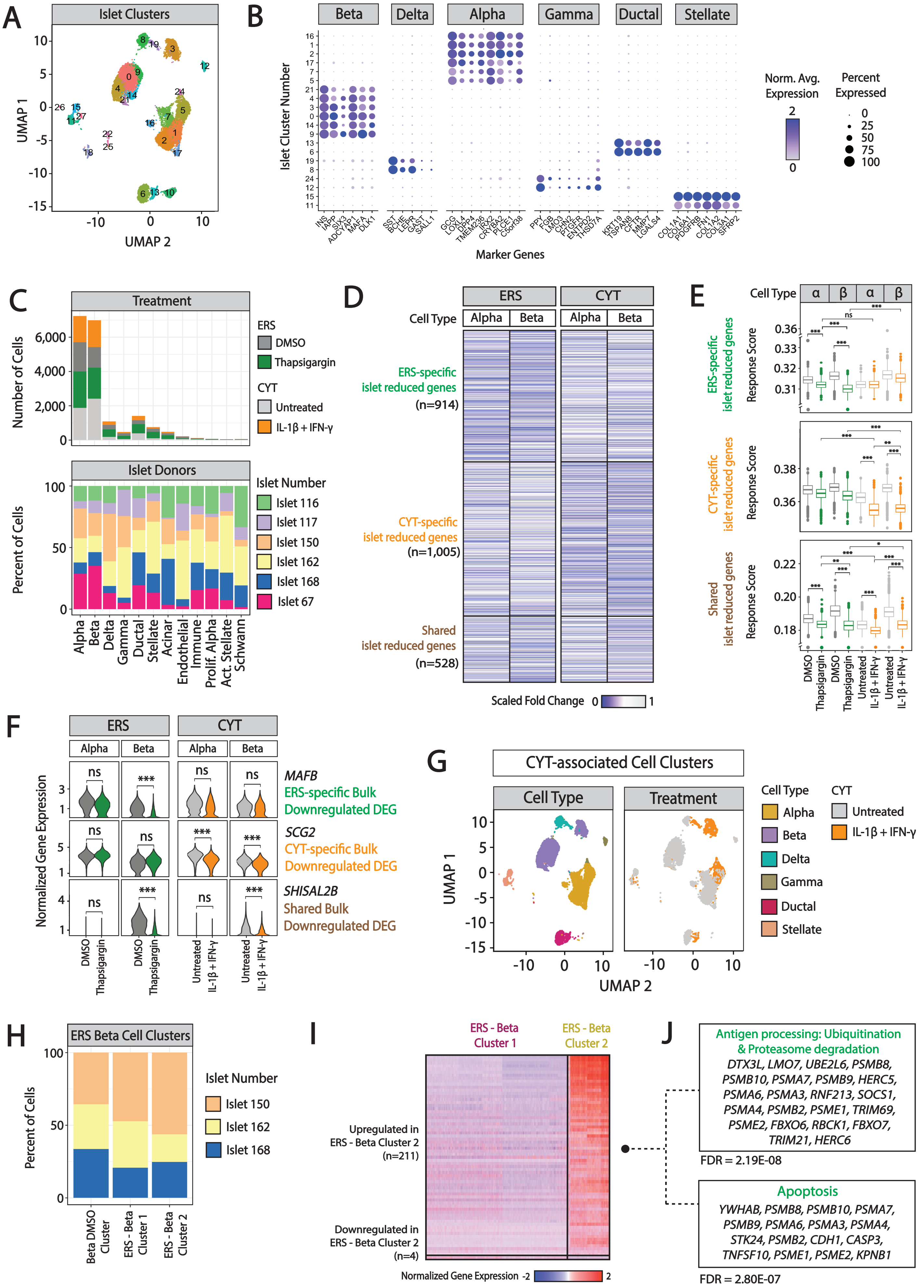

### Supplementary Figure 3

SUPPLEMENTARY FIGURE 3

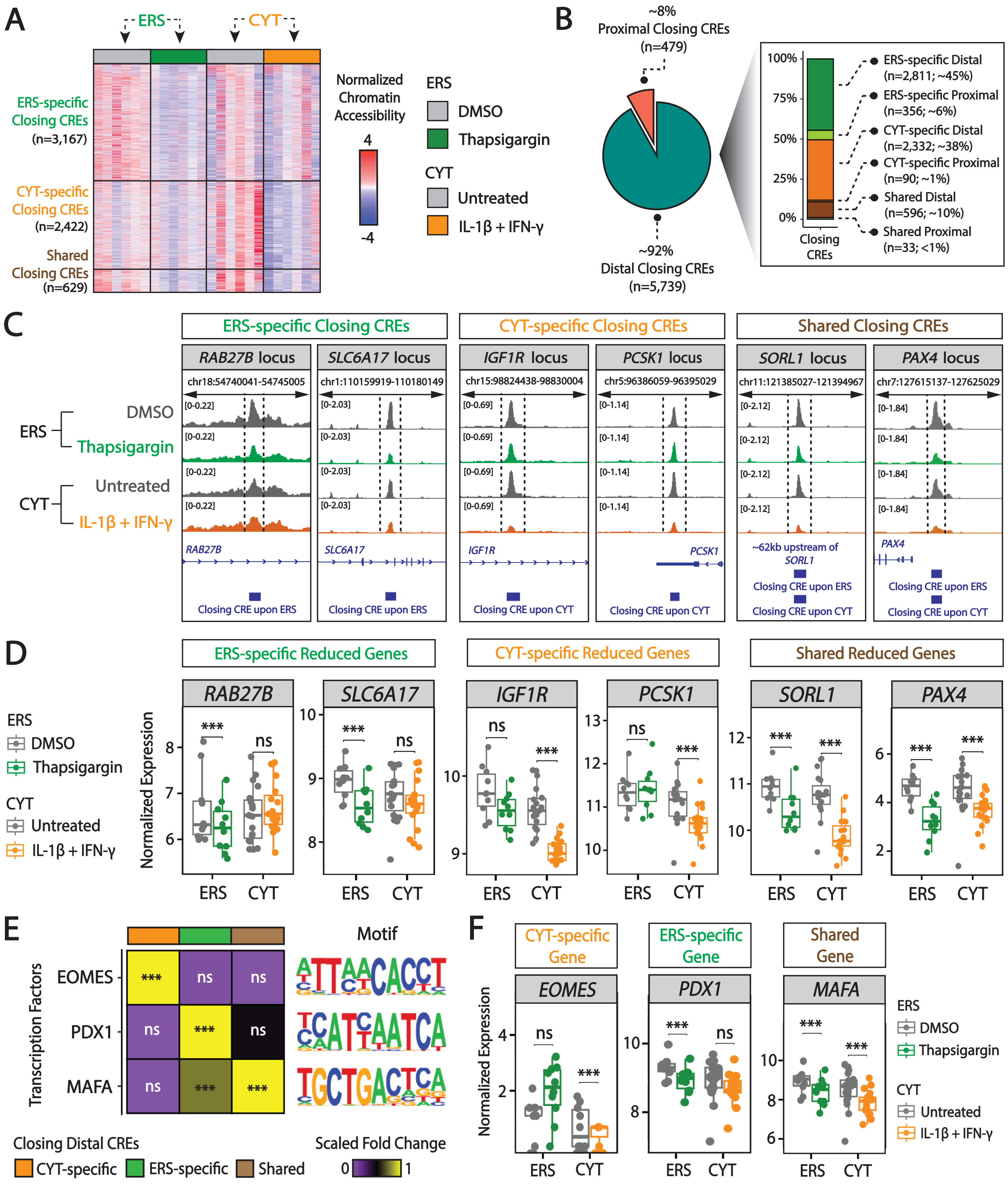

### Supplementary Figure 5

SUPPLEMENTARY FIGURE 5

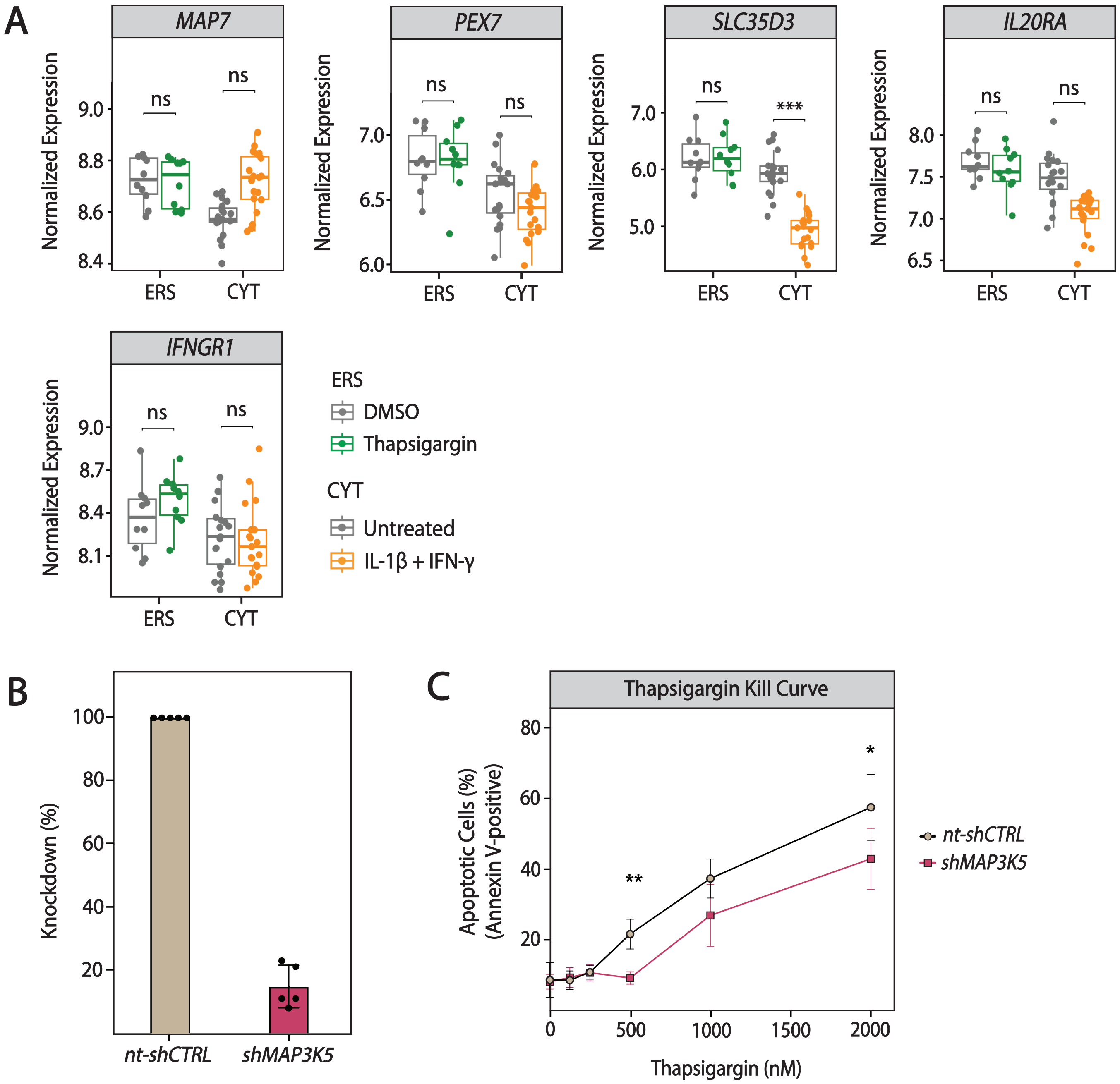
