## Supplementary Figure 4 for "Comparative Multi-omic Mapping of Human Pancreatic Islet Endoplasmic Reticulum and Cytokine Stress Responses Provides Insights into Type 2 Diabetes Genetics"

A

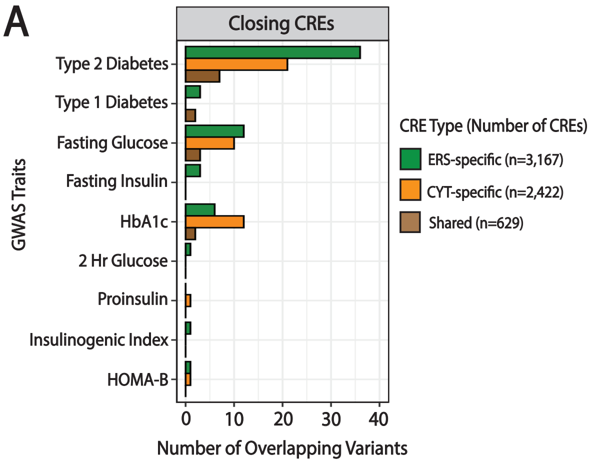


B

**T2D Variants < 500 kb from Induced Gene TSS**

| CYT-specific Opening CRE | CYT-specific Induced Gene | T2D Variant | Alleles 5′→3′ | Risk Allele | Variant-Gene TSS Distance (kb) |
|---|---|---|---|---|---|
| chr16:15206438-15206837 | *NDE1* | rs377264776 | G>C | G | ~436 |
| chr3:15816819-15817218 | *GALNT15* | rs4685264 | G>A | G | ~358 |
| chr9:123027479-123027878 | *OR1L3* | rs631086 | G>A | A | ~352 |
| chr4:88803558-88803957 | *HERC5* | rs10012528 | C>G | G | ~347 |
| | | rs10012624 | C>A | A | ~347 |
| chr7:102695568-102695967 | *UPK3BL2* | rs75518823 | G>T | G | ~158 |
| chr15:90982972-90983371 | *FES* | rs7180016 | G>A | A | ~99 |
| chr6:29845361-29845760 | *HLA-A* | rs60108193 | C>T | C | ~96 |
| | | rs61538976 | A>G | A | ~96 |
| chr1:26126622-26127021 | *SLC30A2* | rs61776594 | G>A | G | ~90 |
| | | rs61776595 | G>T | G | ~90 |
| chr1:46062637-46063036 | *PIK3R3* | rs1768815 | G>A | A | ~23 |
| chr6:131645732-131646131 | *ENPP3* | rs7766061 | G>A | A | ~17 |
| chr6:131619183-131619582 | *ENPP3* | rs10499183 | T>C | C | ~9 |

| Shared Opening CRE | Shared Induced Gene | T2D Variant | Alleles 5′→3′ | Risk Allele | Variant-Gene TSS Distance (kb) |
|---|---|---|---|---|---|
| chr15:62119331-62119736 | *C2CD4B* | rs58728679 | T>- | T | ~44 |

C

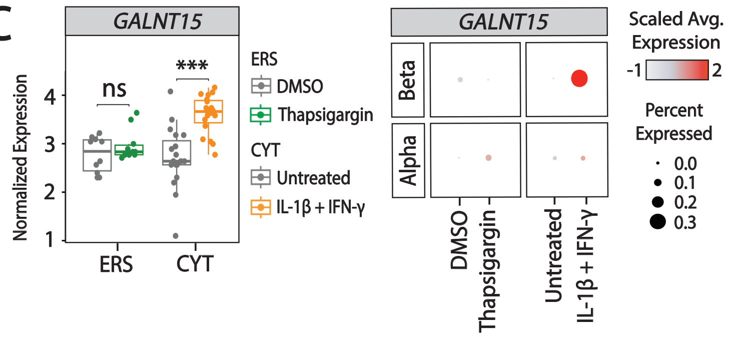


D

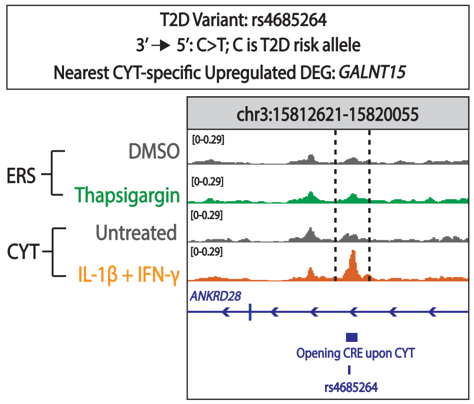


E

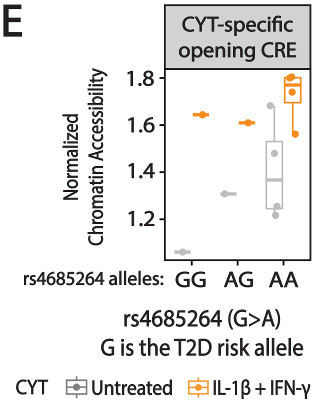


F

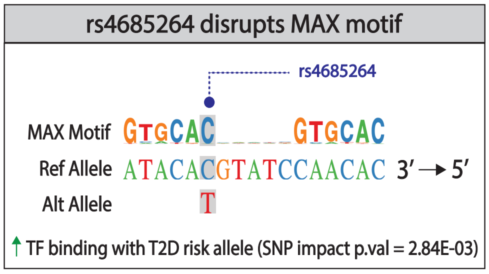


G

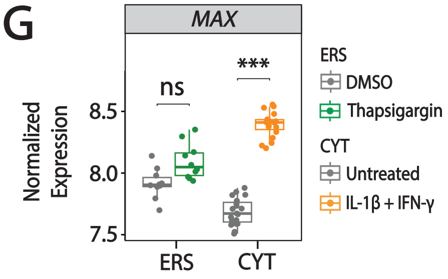
