## Supplementary Figure Legends for "Comparative Multi-omic Mapping of Human Pancreatic Islet Endoplasmic Reticulum and Cytokine Stress Responses Provides Insights into Type 2 Diabetes Genetics"

**Supplementary Figure 1: Reduced transcriptional responses of human pancreatic islets to ER stress (ERS) and pro-inflammatory cytokines (CYT). (A)** Heatmap of genes reduced by ERS and/or CYT treatment (FDR<5%; FC≤0.667). Reduced genes are categorized as ERS-specific, CYT-specific, or shared between both conditions; the number of genes in each category is denoted in parentheses on the left. Note that the majority of genes exhibit stress-specific reduction. Expression values are scaled using z-scores. **(B)** Enriched pathways for reduced genes; FDR values and example genes for each pathway are as indicated. **(C)** Examples of enriched pathway genes reduced by ERS, CYT, or both. Dot-and-box plots show gene expression levels (CPM) per islet donor in ERS (green), CYT (orange), or control samples (grey). ***=FDR<5% and FC≤0.667; ns=not significant. FDRs were calculated using Benjamini-Hochberg p-value adjustment. FDR, False Discovery Rate; FC, fold change; CPM, counts per million.

**Supplementary Figure 2: Single cell resolution of human pancreatic islet responses to ER stress and pro-inflammatory cytokines. (A)** Uniform Manifold Approximation and Projection (UMAP) of aggregated human pancreatic islet single cell transcriptomes (n=3 donors per treatment condition). UMAP is color-coded based on the assigned Seurat cluster. Clusters are numbered for reference. **(B)** Marker gene expression levels across the Seurat cell clusters. Each column corresponds to an islet cell type and the rows represent individual cell clusters. Dot size indicates the percent of cells expressing this gene in each cluster; dot color denotes the scaled average expression level of this gene in those cells. **(C)** Number of cells in each islet cell-type annotated by treatment (top) and by percent contribution from each donor (bottom). **(D)** Scaled fold-change of alpha or beta cell expression of genes reduced by ERS or CYT in whole islets. Genes are grouped into genes whose reduction is ERS-specific, CYT-specific, or shared between conditions. **(E)** Response scores for the islet-reduced genes in alpha and beta cells. ***=p<1.0E-10, **=p<1.0E-05, *=p<1.0E-01; ns=not significant, two-sided Wilcoxon test. **(F)** Violin plots of alpha or beta cell expression for representative genes from the three reduced gene sets in panel D. ***=FDR<5%, FC≤0.667; ns=not significant. **(G)** UMAP visualization of islet scRNA-seq profiles (left) in islets upon CYT (right). **(H)** Percent contribution of each islet donor to ER stress-Beta Cluster 1 (ER stress-BC1), ER stress-Beta Cluster 2 (ER stress-BC2), and beta DMSO control cluster. **(I)** Heatmaps of significantly induced genes in ER stress-BC1 vs. ER stress-BC2 (FDR<5%). Number of induced genes in each category is indicated in parentheses. Expression values are scaled using z-score. **(J)** List of significantly enriched pathways enriched for ER stress-BC2. FDR values for enriched pathways are reported beneath each category. FC, fold-change; $\alpha$, alpha; $\beta$, beta.

**Supplementary Figure 3: Decreased chromatin accessibility changes and associated reduced transcriptional regulatory effects of human islet ER stress (ERS) and pro-inflammatory cytokine (CYT) responses. (A)** Heatmap of human islet *cis*-regulatory elements (CREs) whose accessibility is decreased by ERS and/or CYT treatment (FDR<5%). n=number of CREs in each category. Accessibility values are scaled using z-scores. **(B)** Pie chart showing the percent of closing CREs that are proximal vs. distal (≤1kb vs. >1kb to nearest transcription start site (TSS), respectively). Bar chart (right) indicates the percentage of closing CREs per condition. **(C)** Integrative Genomics Viewer (IGV) browser tracks of representative loci highlighting CREs whose decreased accessibility is ERS-specific, CYT-specific, or shared. **(D)** Dot-and-box plots of gene expression levels (CPM) per islet donor in treated versus control conditions for responsive genes in representative loci in panel C. ***=FDR<5%, FC≤0.667; ns=not significant. **(E)** Heatmap of enriched transcription factor (TF) motifs identified in ERS-specific, CYT-specific, or shared closing distal CREs. The color gradient indicates the scaled fold change of the motif (i.e., motif instances found in target sequences compared to the background sequences). ***=FDR<1.0E-02; ns=not significant. **(F)** Dot-and box plots of islet RNA-seq expression levels (CPM) in ERS, CYT, or control conditions for TF-encoding genes with enriched TF motifs in panel E. ***FDR<5%, FC≤0.667; ns=not significant. False discovery rates (FDR) are calculated using Benjamini-Hochberg p-value adjustment. FC, fold-change; CPM, counts per million.

**Supplementary Figure 4: Type 2 Diabetes (T2D)-associated variants overlapping stress-responsive closing and cytokine (CYT)-specific opening CREs. (A)** Bar chart displaying the number of T2D- or glycemic trait-associated genome-wide association study (GWAS) variants that overlap closing, *cis*-regulatory elements (CREs). **(B)** T2D-associated variants overlapping cytokine (CYT)-specific (top) and shared (bottom) opening CREs located <500 kb from the TSS of a cytokine-specific or shared induced gene, respectively. **(C)** Expression of *GALNT15*, the putative effector gene of T2D variant rs4685264, under ERS and pro-inflammatory cytokine (CYT) conditions in human islet RNA-seq (left) or scRNA-seq (right) profiles. Dot-and-box plots show gene expression levels (CPM) per islet donor in treated versus control samples. ***=FDR<5%, FC≥1.5; ns=not significant. Dot plot of *GALNT15* expression in alpha vs. beta cell scRNA-seq profiles in ERS or CYT treated human islets (right). Dot size indicates the percent of *GALNT15* expressing cells in each cell type; dot color denotes the scaled average *GALNT15* expression in those cells. **(D)** Integrative Genomics Viewer (IGV) browser track showing a CYT-specific opening CRE containing T2D-associated variant rs4685264. **(E)** Dot-and-box plots of islet chromatin accessibility levels (CPM) in donors with rs4685264 GG, AG or AA genotypes (on the plus strand). Note that the homozygous T2D risk allele (G) genotype is associated with the highest *in vivo* chromatin accessibility. **(F)** Composite logo plot (generated using atSNP[164]) illustrates that the rs4685264 T2D risk allele (G on plus strand, C on minus strand) significantly alters a MAX (indicated by the position weight matrix) transcription factor (TF) binding motif (atSNP p-value=2.84E-03) to create a binding site. **(G)** Expression of *MAX*, the gene encoding MAX, in ERS, CYT or control conditions. Dot-and-box plots show gene expression levels (CPM) per islet donor in treated versus control samples. ***=FDR<5%; FC≥1.5; ns=not significant. False discovery rates (FDR) are calculated using Benjamini-Hochberg p-value adjustment. FC, fold-change; CPM, counts per million.

**Supplementary Figure 5: Expression of genes within ±500kb of Type 2 Diabetes (T2D) GWAS variant rs6917676, and the effect of its putative effector gene, *MAP3K5*, on beta cell apoptosis in response to ER stress. (A)** Expression of genes present within the ±500kb window from the T2D variant rs6917676, under ERS and CYT conditions. Dot-and-box plots show gene expression levels (CPM) per islet donor in treated versus control samples. ***=FDR<5%; FC≤0.667; ns=not significant. **(B)** TaqMan assay of *MAP3K5* expression in *nt-shCTRL* vs. *shMAP3K5* EndoC-βH3 cells. **(C)** Thapsigargin kill curve (mean ± standard deviation) representing the percent of apoptotic (Annexin V-positive) cells at varying concentrations of thapsigargin in *MAP3K5* knockdown (*shMAP3K5*, magenta) or non-targeting *shRNA* control (*nt-shCTRL*, black) EndoC-βH3 cells. **=p<0.01; *p=<0.05, two-tailed t-test. False discovery rates (FDR) are calculated using Benjamini-Hochberg p-value adjustment. FC, fold-change; CPM, counts per million.
